# PAX3-FOXO1 coordinates enhancer architecture, eRNA transcription, and RNA polymerase pause release at select gene targets

**DOI:** 10.1101/2021.10.03.462944

**Authors:** Susu Zhang, Jing Wang, Qi Liu, W. Hayes McDonald, Monica L. Bomber, Hillary M. Layden, Jacob Ellis, Scott C. Borinstein, Scott W. Hiebert, Kristy R. Stengel

**Affiliations:** Department of Biochemistry, Vanderbilt University School of Medicine, Nashville, Tennessee 37232, USA; Department of Biostatistics, Vanderbilt University School of Medicine, Nashville, Tennessee 37203, USA; Center for Quantitative Sciences, Vanderbilt University Medical Center, Nashville, Tennessee 37232, USA; Department of Pediatrics, Vanderbilt University School of Medicine, Vanderbilt University Medical Center, Nashville, Tennessee 37203, USA; Vanderbilt-Ingram Cancer Center, Nashville, Tennessee 37027, USA

## Abstract

Transcriptional control is a highly dynamic process that changes rapidly in response to various cellular and extracellular cues^1^. Thus, it is difficult to achieve a mechanistic understanding of transcription factor function using traditional genetic deletion or RNAi methods, because these slow approaches make it challenging to distinguish direct from indirect transcriptional effects. Here, we used a chemical-genetic approach to rapidly degrade a canonical transcriptional activator, PAX3-FOXO1^2-6^ to define how the t(2;13)(q35;q14) disrupts normal gene expression programs to trigger cancer. By coupling rapid protein degradation with the analysis of nascent transcription over short time courses, we identified a core transcriptional network that rapidly collapsed upon PAX3-FOXO1 degradation. Moreover, loss of PAX3-FOXO1 impaired RNA polymerase pause release and transcription elongation at regulated gene targets. The activity of PAX3-FOXO1 at enhancers controlling this core network was surprisingly selective and often only a single element within a complex super-enhancer was affected. In addition, fusion of the endogenous PAX3-FOXO1 with APEX2 identified proteins in close proximity with PAX3-FOXO1, including ARID1A and MYOD1. We found that continued expression of PAX3-FOXO1 was required to maintain chromatin accessibility and allow neighboring DNA binding proteins and chromatin remodeling complexes to associate with this small number of regulated enhancers. Overall, this work provides a detailed mechanism by which PAX3-FOXO1 maintains an oncogenic transcriptional regulatory network.

## Main

The t(2;13) chromosomal translocation is the initiating event in aggressive alveolar rhabdomyosarcoma (aRMS) and fuses the PAX3 DNA binding domain to the C-terminal regulatory domains of FOXO1^2-6^. A thorough understanding of PAX3-FOXO1 function is necessary to both inform aRMS etiology, as well as identify novel therapeutic opportunities. Genetic inactivation coupled with RNA-seq and chromatin immunoprecipitation has suggested that DNA-binding transcription factors can control the expression of thousands of genes, but faster methods of transcription factor inactivation suggest that this is an over estimation due to indirect and compensatory transcriptional changes^7,8^. Small molecule “degraders” coupled with the analysis of nascent transcripts and proteomics offer an opportunity to greatly refine these transcription factor networks and define mechanisms of transcriptional control^9,10^.

### PAX3-FOXO1 degradation impairs RNA pol II pause release at gene targets

We used CRISPR-based genome editing to integrate a 2x*HA-FKBP12*^*F36V*^ degron tag into the last exon of the endogenous *PAX3-FOXO1*^11-13^ locus of Rh30 aRMS cells (Extended Data Fig. 1a). This yielded a model for rapid PAX3-FOXO1 protein degradation following the addition of a small molecule proteolysis-targeting chimera (PROTAC), dTAG-47^14,15^ (Fig. 1a). We generated single cell clones in which the wild-type *FOXO1* locus, the *PAX3-FOXO1* locus, or both were edited. To account for clonal variation, we generated multiple PAX3-FOXO1-2xHA-FKBP12^F36V^-expressing clones (Extended Data Fig. 1b). RNA-seq was performed in the absence of dTAG-47, to demonstrate that the addition of the degron tag did not significantly alter gene expression patterns relative to unedited Rh30 cells (Extended Data Fig. 1c-d). Furthermore, all clones exhibited a similar growth inhibition following PAX3-FOXO1 degradation (Extended Data Fig. 1e). We chose clone 10 for further analysis, and observed altered cell morphology, hallmarks of myogenic differentiation, G_1_ cell cycle arrest, and reduced growth in soft agar following PAX3-FOXO1 degradation (Fig. 1b, Fig. 1c, Extended Data Fig. 2). In contrast, degradation of FOXO1-FKBP12^F36V^ had no effect on cell growth, viability, or gene expression, likely due to the nuclear exclusion of FOXO1 in these cells (Extended Data Fig. 3).

**Figure 1.**
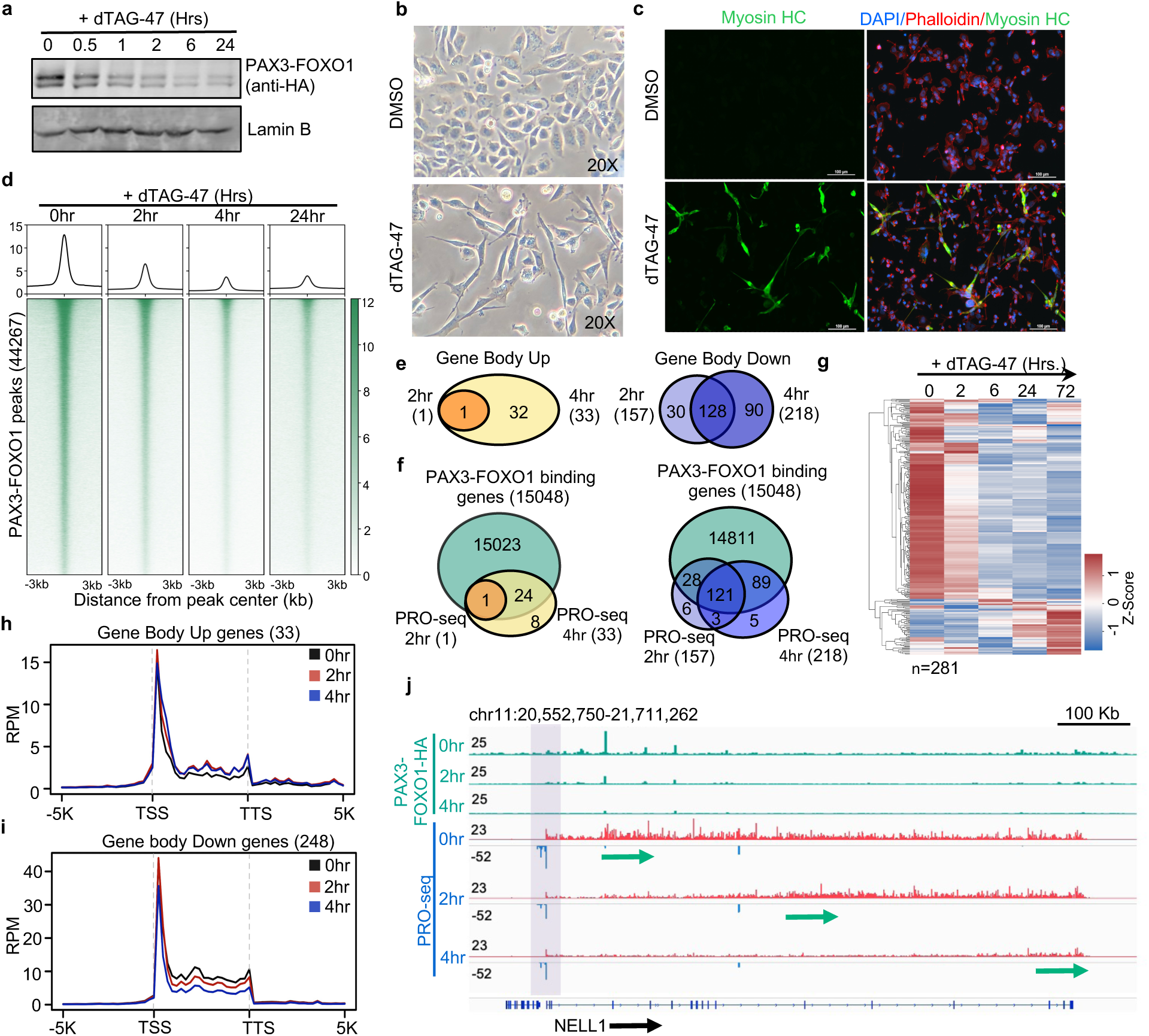
Degradation of PAX3-FOXO1 triggers cell death, differentiation and a loss of expression of a small number of genes. **a**, Western blot analysis showing degradation of endogenous PAX3-FOXO1. **b**, Morphological analysis of PAX3-FOXO1 cells after treatment with dTAG-47 for 6 days (20X). **c**, Immunofluorescence detection of Myosin Heavy Chain positive cells. **d**, Heatmaps of PAX3-FOXO1 CUT&RUN peaks after treatment with dTAG-47. **e & f**, Venn diagrams showing the PRO-seq changes over time (**e**), and the changed genes occupied by PAX3-FOXO1 (**f**). **g**, Heatmap of RNA-seq-detected mRNAs corresponding to genes changed in PRO-seq analysis. **h & i**, Metagene plots of PRO-seq reads of all genes with increased (**h**) or decreased (**i**) transcription. **j**, Genome browser view of *NELL1* showing RNA polymerase moving down the gene (arrows). Box highlights paused polymerase on the sense (red) and anti-sense (blue) strands.

The inclusion of a 2xHA tag in the degron cassette allowed for the robust detection of PAX3-FOXO1 chromatin association by CUT&RUN^16^ (Fig. 1d). A reduction of chromatin-bound PAX3-FOXO1 at nearly all of the over 44,000 identified peaks was observed within 2hr following dTAG-47 treatment, and maximal reduction of chromatin bound PAX3-FOXO1 was observed after 4hr (Fig. 1d). Next, we performed precision nuclear run-on transcription sequencing (PRO-seq) at short time points following PAX3-FOXO1 degradation. PRO-seq maps transcriptionally engaged RNA polymerase across the genome, and provides a readout of nascent transcription, as well as assessments of RNA polymerase pausing and elongation^17,18^. Relative changes in gene body transcription were quantified using the nascent RNA-sequencing analysis (NRSA)^19^ pipeline, which identified only 1 gene with increased transcription at 2hr and only 33 within 4hr of PAX3-FOXO1 degradation (Fig. 1e, left). In contrast, 157 genes exhibited decreased gene body transcription at 2hr following PAX3-FOXO1 degradation, and transcription of most of these genes remained reduced at 4 hours (Fig. 1e, right), which is consistent with the established role for PAX3-FOXO1 in transcriptional activation. Moreover, the vast majority of down-regulated genes were associated with nearby PAX3-FOXO1 CUT&RUN peaks (Fig. 1f; 149 genes at 2hr and 210 at 4hr). Furthermore, the reduced polymerase activity detected by PRO-seq resulted in a reduction of mature mRNA as detected by RNA-seq (Fig. 1g), and was associated with reduced H3K4me3 at regulated promoters (Extended Data Fig. 4a), nominating these genes as direct PAX3-FOXO1 gene targets. It is notable that several transcription factors were turned off by degradation of PAX3-FOXO1, including oncogenes and “stemness” factors (e.g., *JUN, KLF4, RUNX2, ETS1, PRDM12* and the co-repressor *RUNX1T1*, Table 1), emphasizing the need to use nascent transcript analysis at early timepoints in order to avoid detection of secondary transcriptional changes found by RNA-seq at later times (Extended Data Fig. 4b).

Next, we generated histograms depicting the PRO-seq signal across genes up-regulated (33, Fig. 1h) or down-regulated (248, Fig. 1i) in at least one timepoint following PAX3-FOXO1 degradation. Interestingly, down-regulated genes exhibited a clear reduction in gene body polymerase density, but maintained high levels of paused polymerase just downstream of the transcription start site (Fig. 1j). This pattern is indicative of a reduction in RNA polymerase pause release rather than a significant change in transcription initiation. In fact, most down-regulated genes showed an increase in RNA polymerase pausing index, which reached statistical significance at 106 of 248 down-regulated genes (Extended Data Fig. 4c-d, Table 1). Notably, those genes with a significant increase in pausing index were more highly down-regulated than genes without changes in pausing index (Extended Data Fig. 4e). Examination of long genes such as *NELL1* further illustrates this phenomenon (Fig. 1j). By 2hr following PAX3-FOXO1 degradation, there was a gap between the paused polymerase and elongating RNA polymerase moving through the gene body, and by 4hr the elongating polymerase had reached the 3’ end of the gene (Fig. 1j, arrows). During this time, levels of paused polymerase were only slightly reduced for both sense and antisense transcripts (Fig. 1j, shaded box). Thus, PAX3-FOXO1 regulated transcription of many target genes by promoting RNA pol II pause release and transcription elongation.

Greater than 80% of PAX3-FOXO1 CUT&RUN peaks were localized to intergenic or intronic regions (Extended Data Fig. 4f), suggesting that PAX3-FOXO1 functions at enhancers to regulate gene expression. PRO-seq detects all nascent transcripts, therefore, NRSA was used to quantify intergenic enhancer RNA (eRNA) transcription following PAX3-FOXO1 degradation (Fig. 2a). Within the first 4hr of PAX3-FOXO1 degradation, 305 eRNAs were significantly down-regulated and 289 (96%) of these overlapped with PAX3-FOXO1 binding sites identified by CUT&RUN (Fig. 2b). Given this high overlap, we performed ChIP-seq for histone H3K27ac and CUT&RUN for BRD4, which are hallmarks of active enhancers^20^. Globally, there were very few changes in either H3K27ac or BRD4 occupancy following PAX3-FOXO1 degradation (Extended Data Fig. 4g-h). We then compared changes in H3K27ac and BRD4 occupancy between all PAX3-FOXO1 peaks (44,267, Fig. 2c, left) and PAX3-FOXO1 peaks associated with changes in eRNA production following PAX3-FOXO1 degradation (289, Fig. 2c, right). While a similar reduction of PAX3-FOXO1 was observed at both sets of PAX3-FOXO1 peaks (Fig. 2c), there was a robust reduction in H3K27ac and BRD4 levels only at the 289 PAX3-FOXO1 peaks associated with changes in eRNA synthesis (Fig. 2c, right panels). It is notable that this number likely underestimates the number of enhancers regulated by PAX3-FOXO1, because it does not include intronic enhancers that confound eRNA analysis due to gene body transcription^19^. Nevertheless, this suggests that while PAX3-FOXO1 associates with chromatin at many sites throughout the genome, only a small number of those sites were associated with changes in enhancer function following PAX3-FOXO1 degradation.

**Figure 2.**
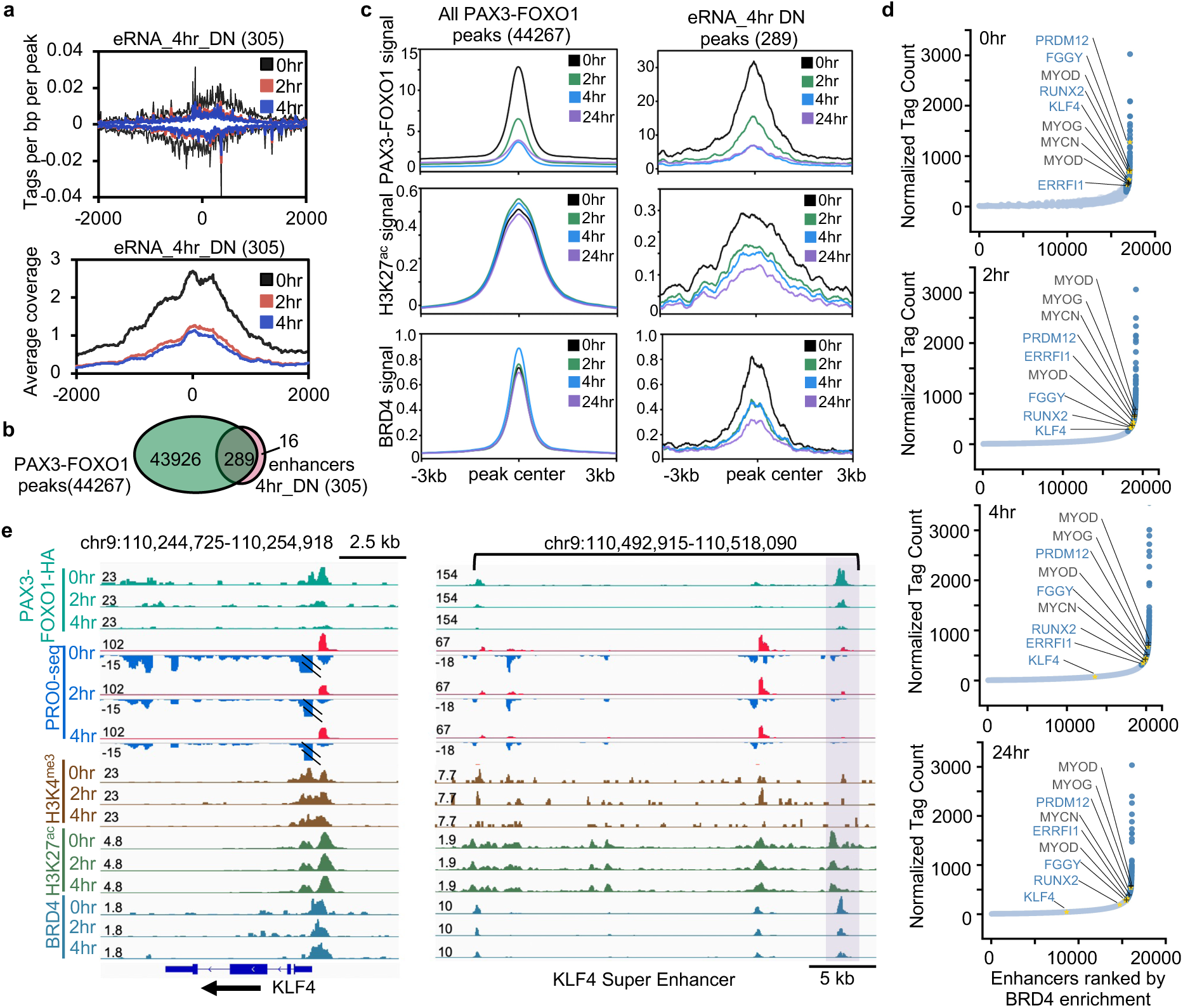
PAX3-FOXO1 maintains active enhancers. **a**, Histograms of eRNA signals after PAX3-FOXO1 degradation. Upper panel, tags per base pair; lower panel, average read count. **b**, Venn diagram of overlap between PAX3-FOXO1 peaks and down-regulated eRNAs. **c**, Average signal of PAX3-FOXO1, H3K27ac, and BRD4 after dTAG-47 treatment over all PAX3-FOXO1 bound sites vs. PAX3-FOXO1 sites showing eRNA changes. **d**, Super-enhancers (dark blue) were identified using BRD4 binding before and after PAX3-FOXO1 degradation. Gold asterisks, super-enhancers regulated by PAX3-FOXO1; black +, previously identified PAX3-FOXO1-associated super-enhancers not changed upon PAX3-FOXO1 degradation. **e**, IGV gene tracks showing a time course of dTAG-47 treatment for PRO-seq and genomic localization of factors and histone marks. Shaded area, PAX3-FOXO1 peak.

PAX3-FOXO1 has been suggested to establish super-enhancers to drive myogenic transcription networks^21-23^. Therefore, we identified super-enhancers based on BRD4 enrichment^20,24-26^ and asked whether continued PAX3-FOXO1 expression was required to maintain these regulatory structures (Fig. 2d). Our analysis identified many of the super-enhancers previously associated with PAX3-FOXO1 function, but degradation of PAX3-FOXO1 did not affect the super-enhancers of *MYOD1, MYOG*, and *MYCN*^6^. Moreover, even though all three of these super-enhancers were bound by PAX3-FOXO1 (Extended Data Fig. 5a-c), we detected no change in *MYCN* transcription, an increase in *MYOG* transcription, and only a minimal decrease in *MYOD1* transcription at 4 hours, which did not result in altered *MYOD1* mRNA levels by RNA-seq (Fig. 1, 2).. Moreover, the *MYOD1, MYOG* and *MYCN* super-enhancers persisted even 24 hours after PAX3-FOXO1 degradation (Fig. 2d), suggesting that PAX3-FOXO1 was not required for their maintenance. In contrast, we identified super-enhancer associated genes that were rapidly down-regulated following PAX3-FOXO1 degradation, including *RUNX2, KLF4, FGGY*, and *PRDM12* (Fig. 2d). Interestingly, these super-enhancers fell lower on the BRD4-defined super-enhancer list over time. Manual inspection of each of these super-enhancers showed that they were bound by PAX3-FOXO1, but rather than a complete collapse of the super-enhancer, there was a disruption of selective PAX3-FOXO1-bound enhancer elements upon degradation of PAX3-FOXO1 (Fig. 2e). For instance, the region upstream of *KLF4* and the intronic enhancers of *FGGY* contained multiple enhancers marked by eRNA production, H3K27ac, and BRD4 binding. However, only one of the enhancer elements (box, Fig. 2e for *KLF4* and Extended Data Fig. 5d for *FGGY*) showed a rapid reduction in H3K27ac and BRD4 binding upon PAX3-FOXO1 degradation. Thus, disruption of a single enhancer element within the enhancer cluster was associated with a significant reduction in gene body transcription (Fig. 2e, left, PRO-seq panels).

### Proteomics analyses reveal PAX3-FOXO1-associated protein complexes

We next sought to define protein complexes that contribute to PAX3-FOXO1-mediated transcription activation. Previous proteomic analysis of PAX3-FOXO1 relied on over-expression of the fusion protein^27^. Therefore, we modified our CRISPR homology-directed repair vector to generate an endogenous PAX3-FOXO1-APEX2 protein fusion. APEX2 is an engineered peroxidase, which creates biotin-phenoxyl radicals that covalently bind nearby proteins (<20 nm)^28-30^. Thus, proteins in close proximity to PAX3-FOXO1 were purified using streptavidin beads and identified by liquid chromatography coupled with mass spectrometry. We identified over 500 significantly enriched proteins including components of multiple transcriptional complexes, such as FACT and SWI/SNF, transcription elongation factors (e.g. CDK9, CCNT1, NELFB), and components of a Mediator subcomplex that is associated with transcriptional elongation^31^ (MED12 and CDK8, Fig. 3a-b). Interestingly, while we did not identify an association with BRD4^6^, we did identify the enzymes that place the histone mark bound by BRD4 (TRRAP and EP400)^32^. We also identified other sequence-specific transcription factors, which may cooperate with PAX3-FOXO1 to regulate gene expression, including MYOD, HEB (TCF12), and RUNX1/CBFB (Fig. 3a-b). Motif analysis of PAX3-FOXO1 CUT&RUN peaks identified an enrichment of these same transcription factor-binding motifs at PAX3-FOXO1-bound enhancers (Extended Data Fig. 6a), further suggesting that PAX3-FOXO1 regulates gene expression in concert with these factors. Finally, we used CRISPR/Cas9 to integrate a 3xFLAG tag into the endogenous *PAX3-FOXO1* locus and confirmed a number of these associations by affinity purification using the 3xFLAG tag and LC-MS (Extended Data Fig. 6b-c).

**Figure 3.**
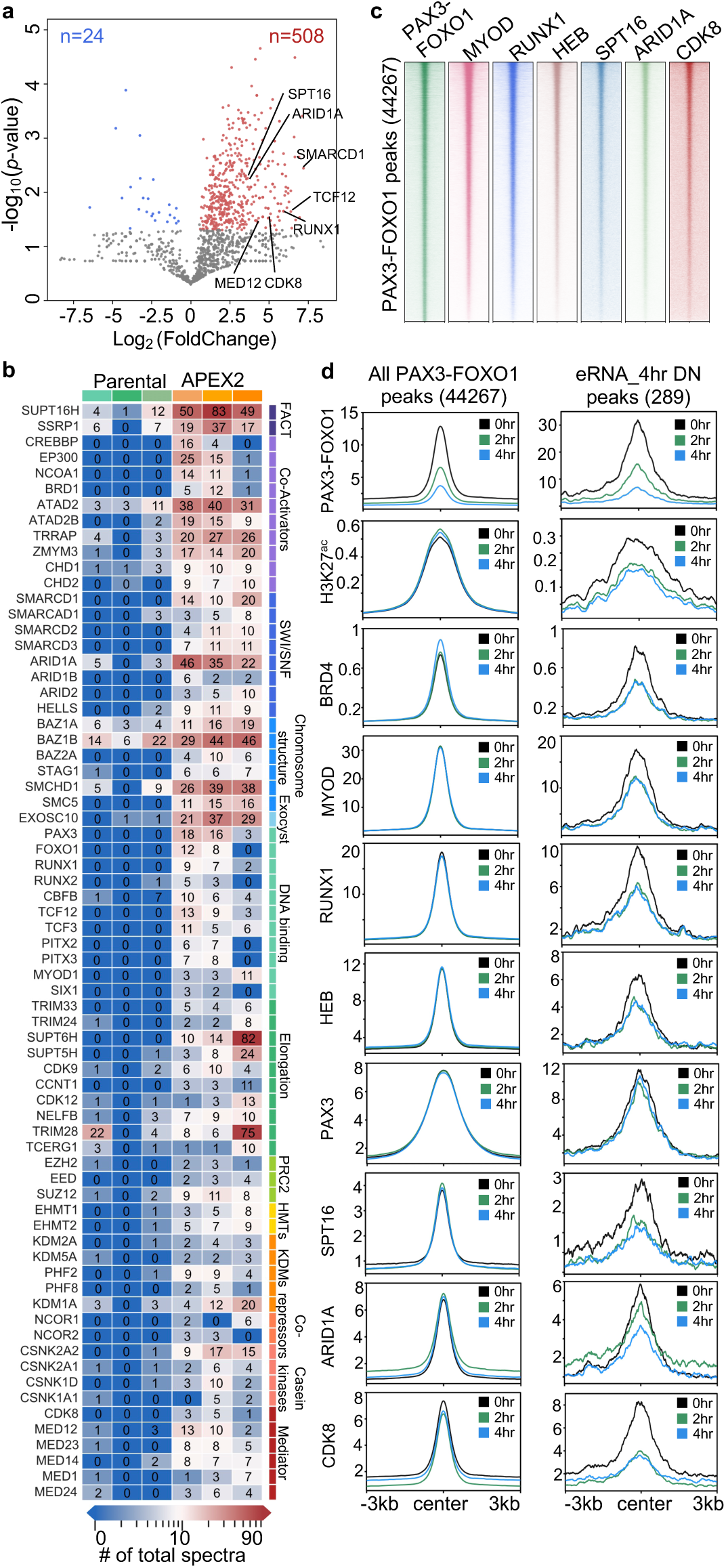
PAX3-FOXO1 recruits complexes involved in transcription. **a**, Volcano plot showing log2-fold change (PAX3-FOXO1-APEX2/Parental) vs. -log10 of the p-value generated from the mass spectrometry results of PAX3-FOXO1-APEX2-mediated biotinylation (n=3, one-tail unpaired T-test). Blue dots, enriched in parental samples; red dots, enriched in PAX3-FOXO1-APEX2 samples. **b**, Heatmaps of selected PAX3-FOXO1-APEX2 biotinylated proteins from the APEX2-mass spectrometry analysis. Total spectral count is shown within each box. **c**, Heatmaps of CUT&RUN analysis of representative factors from the analysis in b plotted around all PAX3-FOXO1 binding sites. **d**, Histograms of the CUT&RUN signal of the indicated factors around all PAX3-FOXO1 peaks and the down-regulated eRNA peaks. Graphs of PAX3-FOXO1, H3K27ac, and BRD4 from Fig. 2c are shown for comparison.

We then performed CUT&RUN for MYOD1, HEB (TCF12), ARID1A (a SWI/SNF component), SPT16 (FACT component), and CDK8 (Mediator component) before and after PAX3-FOXO1 degradation to determine if the APEX2-identified complexes co-localized with PAX3-FOXO1 on chromatin. There was an enrichment of these factors at all PAX3-FOXO1 binding sites (Fig. 3c, Extended Data Fig. 6d-e). While we did not see a global reduction of any of these factors from PAX3-FOXO1 binding sites following PAX3-FOXO1 degradation, we did observe the synchronous loss of all factors from specific PAX3-FOXO1 binding sites associated with regulated eRNAs (Fig. 3d, Extended Data Fig. 6e).

### PAX3-FOXO1 is required to maintain open chromatin at regulated enhancers

The fact that MYOD, RUNX1, HEB, ARID1A, SPT16, and CDK8 were all lost from regulatory PAX3-FOXO1 binding sites within 2hr of PAX3-FOXO1 degradation (Fig. 3d) was surprising and may suggest that PAX3-FOXO1 binding is required to maintain enhancer architecture to allow transcriptional complexes to assemble on chromatin. Therefore, we performed an ATAC-seq^33^ time course following PAX3-FOXO1 degradation. There was a high degree of overlap between PAX3-FOXO1 binding and ATAC-seq-defined nucleosome free regions (NFRs, Extended Data Fig. 7a). While most ATAC-seq signals remained unchanged following PAX3-FOXO1 degradation (Fig. 4a, Extended Data Fig. 7b), there was a significant reduction in accessibility at 810 ATAC-seq-identified NFRs within 2hr of PAX3-FOXO1 degradation (Fig. 4a, Extended Data Fig. 7c) with few increases in accessibility. The number of NFRs with a significant decrease in accessibility grew to 1,226 by 4hr, and by 24hr, there were significant increases in the number of sites exhibiting both increases and decreases in accessibility, suggesting secondary changes in chromatin structure by 24hr (Fig. 4a).

**Figure 4.**
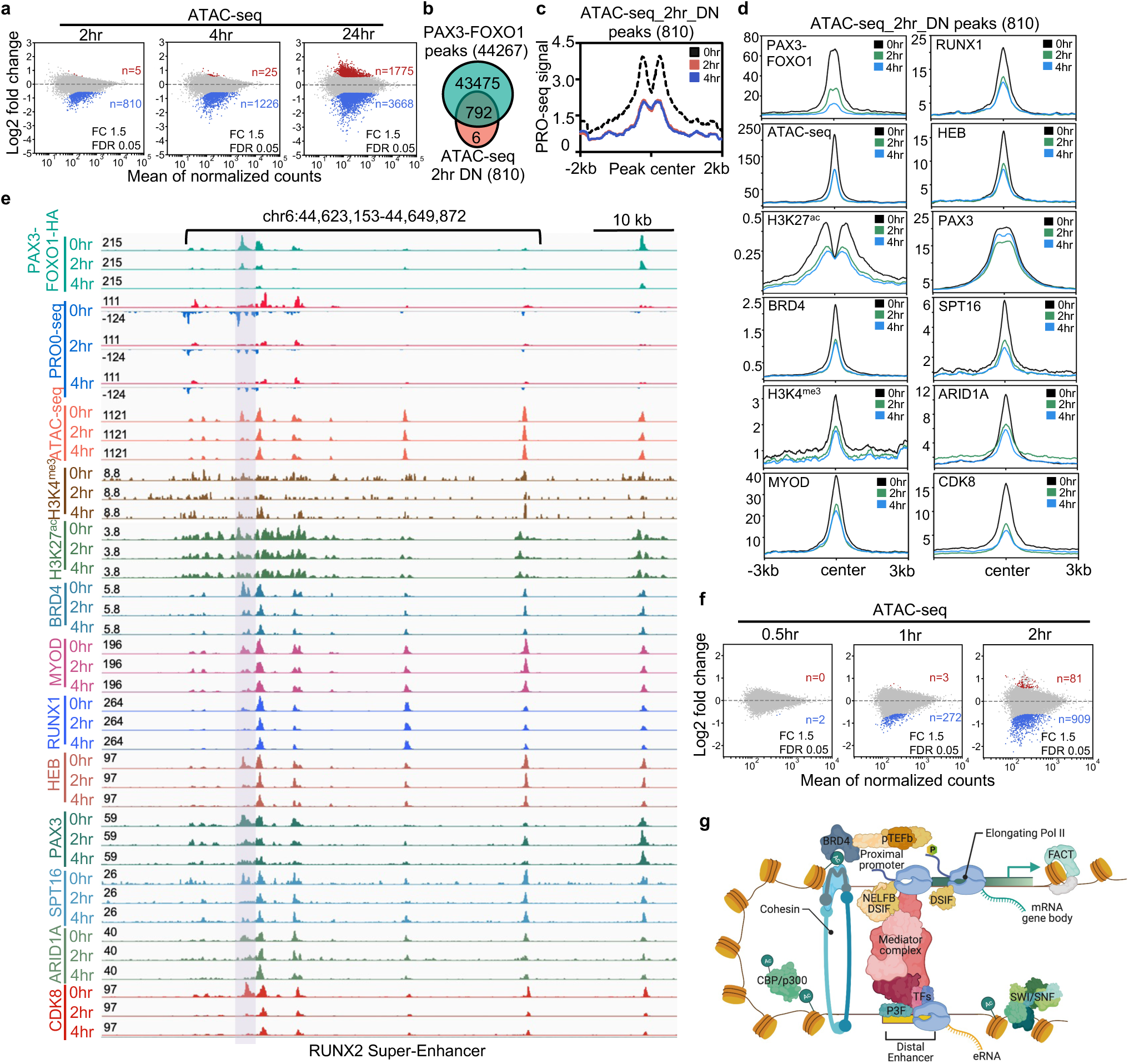
PAX3-FOXO1 is required for maintaining open chromatin structure. **a**, MA plots showing ATAC-seq peak changes; red (up) or blue (down) (dTAG-47/DMSO, n = 2). **b**, Venn diagram of overlap between PAX3-FOXO1 peaks and the significantly down-regulated ATAC-seq peaks. **c**, Histogram of the PRO-seq signal around the significantly down-regulated ATAC-seq peaks. **d**, Histograms of the CUT&RUN signal of the indicated factors around the significantly down-regulated ATAC-seq. **e**, Genome browser view of the *RUNX2* “super” enhancer. The grey shaded box highlights an enhancer showing changes in factor binding, eRNAs, and accessibility over time compared to neighboring enhancers. **f**, MA-plots of ATAC-seq peak changes from 0.5hr to 2hr after dTAG-47 addition. **g**, Model integrating the proteomic and genomic data into a hypothetical model by which PAX3-FOXO1 regulates transcription of its target genes. Created with BioRender.com

Most of the 810 NFRs exhibiting reduced accessibility within 2 hours were located at intergenic and intronic regions bound by PAX3-FOXO1 (Fig. 4b, Extended Data Fig. 7d) and showed a strong enrichment for PAX3-FOXO1 motifs compared with total PAX3-FOXO1-bound loci (compare Extended Data Fig. 6a and 7e). We then associated down-regulated genes with the nearest significantly decreased ATAC-seq or eRNA signal, to define high probability PAX3-FOXO1-regulated enhancer-promoter pairs (Table 2). In particular, 147 down-regulated genes had a nearby reduction in ATAC-seq signal, thus we consider these 147 genes to be high-confidence PAX3-FOXO1 gene targets (Table 2). In addition, there was significant overlap between reduced NFRs and down-regulated eRNAs (Extended Data Fig. 7f, Table 2), and eRNA transcription was significantly reduced at sites of reduced accessibility within 2hr of PAX3-FOXO1 degradation (Fig. 4c). The reverse was also true, as the ATAC-seq signal was specifically reduced when plotted around enhancers that displayed reductions in eRNA synthesis following PAX3-FOXO1 degradation (Extended Data Fig. 7g). When we focused the CUT&RUN analysis of MYOD1, RUNX1, HEB, ARID1A, SPT16, and CDK8 at the 810 changed ATAC-seq peaks, we found a reduction in the signal of each of these factors within 2hr of PAX3-FOXO1 degradation (Fig. 4d, Extended Data Fig. 7h). The loss of MYOD1, RUNX1, HEB, ARID1A, SPT16, and CDK8 binding (Fig. 4d), suggests that PAX3-FOXO1 maintained open chromatin to allow the binding of these factors at specific enhancer elements. The *RUNX2* super-enhancer is exemplary of this theme as only one of the enhancers (grey box, Fig. 4e) was temporally regulated by degradation of PAX3-FOXO1. This enhancer element was characterized by rapid loss of chromatin accessibility, eRNA production, and binding of BRD4, MYOD1, HEB, ARID1A, SPT16, and CDK8. Similar effects were observed at other enhancers, including the *KLF4* and *PRDM12* super-enhancers (Extended Data Fig. 8 and 9). Thus, continued expression of PAX3-FOXO1 was only required to maintain a highly selective set of enhancer elements.

Given that the fusion protein binds to the same consensus motif as PAX3, we next asked whether degradation of PAX3-FOXO1 affected DNA binding by PAX3. CUT&RUN analysis using an antibody to the C-terminus of PAX3 showed that PAX3 bound to the same sites as the fusion protein, including those enhancers that showed a decrease in accessibility following PAX3-FOXO1 degradation (Fig. 4d-e, Extended Data Fig. 7i), suggesting that unlike PAX3-FOXO1, PAX3 was not sufficient to maintain chromatin accessibility at these enhancers. In fact, PAX3 binding was reduced over time following PAX3-FOXO1 degradation much like MYOD and HEB (Fig. 4d-e).

We were surprised to note that enhancer accessibility was so quickly affected following dTAG-47 treatment. Therefore, we performed an even shorter ATAC-seq and PRO-seq time course to define how quickly loss of PAX3-FOXO1 affects chromatin accessibility and enhancer activation (eRNA production). While no changes in chromatin accessibility were observed 30 min following dTAG-47 treatment, by 1hr chromatin accessibility was reduced at 272 sites and by 2hr changes mirrored our previous ATAC experiment (909 vs. 810, compare Fig. 4a and 4f). In fact, when the sites affected at 2hr were plotted as a heat map (Extended Data Fig. 7j), one can see that many of the 909 down-regulated peaks were losing accessibility within the first 30min, but had not reached our significance cut offs (Fig. 4f). Similar to the ATAC-seq analysis, PRO-seq analysis identified few changes in gene body transcription in the first 30min after dTAG-47 treatment, but by 1hr, significant changes in gene body transcription were observed (Extended Data Fig. 7k-m). Combined, the proteomic and genomic data indicate that chromatin was rapidly remodeled following PAX3-FOXO1 degradation, leading to the synchronous loss of transcriptional complexes from a small number of discrete enhancer elements, and ultimately resulting in the loss of transcriptional elongation at PAX3-FOXO1 target genes (Fig. 4g).

## Discussion

Given that transcriptional changes occur rapidly, traditional genetic approaches have failed to effectively define direct targets of sequence-specific transcription factors, and therefore, have inadequately defined mechanisms of transcriptional control by these proteins^9,10^. Thus, CRISPR-based addition of degron tags to endogenous transcription factor proteins has provided a technological breakthrough that is greatly aiding the study of transcription factor function^7,8,34,35^. Here, we have applied this approach to an oncogenic fusion transcription factor, PAX3-FOXO1. While we identified greater than 40,000 PAX3-FOXO1 binding sites throughout the genome, by combining rapid protein degradation with nascent transcript analysis by PRO-seq and enhancer accessibility by ATAC-seq, we determined that PAX3-FOXO1 only activates transcription of approximately 147 high-confidence gene targets. Moreover, we find that many of these targets are regulated at the level of RNA polymerase pause release rather than at the stage of transcription initiation, which defines a new mechanism of PAX3-FOXO1 action.

Previous studies have postulated that PAX3-FOXO1 possesses pioneer activity, facilitating the establishment of *de novo* enhancer elements that drive myogenic transcriptional programs^6,23^. Indeed, we observed the rapid loss of chromatin accessibility at regulated enhancers following PAX3-FOXO1 degradation. Furthermore, we also detected the binding of PAX3-FOXO1 at super-enhancers (Fig. 2d). However, we did not observe the broad collapse of these super-enhancers following PAX3-FOXO1 degradation, but rather a rapid loss of accessibility at specific enhancer elements within the enhancer cluster. Surprisingly, disruption of a single enhancer at PAX3-FOXO1-bound super-enhancers resulted in a significant loss of gene transcription. These studies raise questions as to how individual elements within enhancer clusters cooperate to regulate gene expression and whether some elements act in a dominant fashion, nucleating the overall enhancer activity, or if a single enhancer in a cluster is sufficient to direct transcription.

By engineering the endogenous PAX3-FOXO1 for proximity labeling, we identified DNA binding factors and transcriptional complexes that associate with PAX3-FOXO1 (Fig. 3). Among the identified interacting proteins was ARID1A, which could suggest that the continued recruitment of SWI/SNF may be required to maintain the NFR at enhancers to allow full transcriptional complex assembly. Our data also suggests that PAX3, which was bound at these same loci, is not sufficient to maintain open chromatin at these regulatory elements. Therefore, it seems likely that PAX3-FOXO1, and not PAX3, possesses pioneer activity; however, future studies are needed to more rigorously test this hypothesis. Finally, these data indicate that PAX3-FOXO1 is continuously required to maintain the expression of genes critical for blocking terminal differentiation and to maintain cell viability such as *FGF8*^36^, *RUNX2*^37^, *KLF4*^38^, *PRDM12*^39^, *ALK*, and *FGFR2*^40^ (Table 1) and further emphasize the utility of PAX3-FOXO1 as a therapeutic target in rhabdomyosarcoma.

## Acknowledgements

We especially thank members of the Hiebert lab for helpful discussions, reagents and advice. We thank the Flow Cytometry, Chemical Synthesis, and Genome Sciences (VANTAGE) Shared Resources for services and support. This work was supported by a St. Baldrick’s Research Grant with generous support from Rachael Chaffin’s Research Fund, the T. J. Martell Foundation, the Robert J. Kleberg, Jr. and Helen C. Kleberg Foundation, National Institutes of Health grants (RO1-CA164605, R01-CA255446-01A1, T32-CA009582-33), and as well as core services performed through Vanderbilt Digestive Disease Research grant (NIDDK P30DK58404), the Vanderbilt-Ingram Cancer Center support grant (NCI P30CA68485), and a grant from the National Center for Advancing Translational Sciences (2 UL1 TR000445-06). The content is solely the responsibility of the authors and does not necessarily represent the official views of the NIH.

## Author Contributions

Conceptualization, S.Z., S.W.H and K.R.S.; Methodology, M.L.B., H.M.L., S.Z., J.E., K.R.S.; Investigation, M.L.B, H.M.L, J.E., S.Z., K.R.S.; Formal Analysis, S.Z., J.W., Q.L., and K.R.S.; Writing-Original Draft, S.Z., K.R.S. and S.W.H.; Writing-Review & Editing, K.R.S., J.W., Q.L., S.C.B. and S.W.H.; Visualization, J.W., S.Z., K.R.S. and S.W.H.; Supervision, K.R.S. and S.W.H.; Funding Acquisition, S.W.H. and K.R.S.

## Materials and Methods

### Cell lines

The Rh30 cell line was obtained from ATCC. Cells were cultured in RPMI-1640 (Corning by Mediatech, Inc.) containing 10% FetalPlex (Corning by Mediatech, Inc.) and supplemented with 1% L-Glutamine (Corning by Mediatech, Inc.) and 1% penicillin/streptomycin (Corning by Mediatech, Inc.). *Drosophila* S2 cells (Schneider media supplemented with 10% FBS, 1% penicillin and streptomycin) were a gift from Dr. Emily Hodges.

### dTAG-47

dTAG-47 was synthesized by the Vanderbilt University Medical Center (VUMC) Molecular Design and Synthesis Center (VICB, kk-25-065) as described^15^, and reconstituted in DMSO (Sigma).

### Generation of endogenous PAX3-FOXO1-tagged Rh30 cell lines

The endogenous allele of PAX3-FOXO1 in Rh30 cells was engineered to express C-terminal FKBP12^F36V^-2xHA, APEX2-2xHA, or 3X-FLAG tags using homology-directed DNA repair^41^. 180 bp upstream of the stop codon and 500 bp after the *FOXO1* top codon were cloned into pUC19 containing FKBP12^F36V^-2xHA-P2A-mCherry derived from Addgene #104370 (pAW62^14^) using the Gibson Assembly Cloning Kit (NEB #E5510S). The APEX2 sequence (Addgene #97421) or a 3xFLAG tag was cloned into the HDR donor plasmids to create the *FOXO1-APEX2* and *FOXO1-3xFLAG* plasmids respectively. Cas9, gRNA and the HDR template plasmid were delivered into Rh30 cells by electroporation. mCherry positive cells were sorted and single cell cloning was performed to generate PAX3-FOXO1-tagged and FOXO1-tagged clones.

Primers used to construct template plasmid are listed below:

Primers to generate 5’ homology gene block:

F:GCCAAGTGGGTTGATGTCTGGTTTTTCCTTGAGAGAAGCTCCCAAGTGACTTGGATGGCATGTTC

R:GGGGAGATGGTTTCCACCTGCACTCCTCCGGATCCGCCTGACACCCAGCTATGTGTCGTTGTCTTG

Primers to generate 3’homology gene block:

Cherry-F:

ACTCCACCGGCGGCATGGACGAGCTGTACAAGTAAGGGTTAGTGAGCAGGTAAGTTCACCCCAAT

BFP-F:

ACCTCCCTAGCAAACTGGGGCACAAGCTTAATTAAGGGTTAGTGAGCAGGTAAGTTCACCCCAAT

R:CGGCCAGTGAATTCGAGCTCGGTACCCGGGGATCCCCAAGAAAACTAAAAGGGAGTTGGTGAAAG

5’Gibson Cloning Primers:

F:CAAGACAACGACACATAGCTGGGTGTCAGGCGGATCCGGAGGAGTGCAGGTGGAAACCATCTCCCC

R:GAACATGCCATCCAAGTCACTTGGGAGCTTCTCTCAAGGAAAAACCAGACATCAACCCACTTGGC

3’Gibson Cloning Primers:

Forward:

CTTTCACCAACTCCCTTTTAGTTTTCTTGGGGATCCCCGGGTACCGAGCTCGAATTCACTGGCCG

Cherry-R:

ATTGGGGTGAACTTACCTGCTCACTAACCCTTACTTGTACAGCTCGTCCATGCCGCCGGTGGAGT

BFP-R:

ATTGGGGTGAACTTACCTGCTCACTAACCCTTAATTAAGCTTGTGCCCCAGTTTGCTAGGGAGGT

FOXO1 crRNA: 5’-CAGGCTGAGGGTTAGTGAGC

### Western Blot

Cells were collected and lysed with RIPA buffer (50 mM Tris pH8.0, 150 mM NaCl, 1% NP-40, 0.5% sodium deoxycholate, 0.1% SDS) containing protease inhibitor. After sonication, lysates were cleared by centrifugation and subjected to SDS-PAGE and electrophoretic transfer to membranes before incubation with antibodies directed against HA (Abcam, ab18181), FLAG (Sigma, F1804), GAPDH (Santa Cruz, sc-365062), Lamin B (Santa Cruz, sc-6217). Signal was visualized with secondary IR-Dye conjugated antibodies (Licor) and detected using the Licor Odyssey imaging system.

### Cell growth

Cells were seeded at a density of 2 × 10^5^ cells/ml on day 0 in 6-well culture plates and treated with DMSO or 500 nM dTAG-47. The cells were reseeded every 3 days at 2 × 10^5^ cells/ml and maintained in DMSO/dTAG-47 treatment for the duration of the assay. Viable cell were counted with Trypan Blue dye exclusion every day for 9 days consecutive. Quantification was performed in triplicate and the values averaged and shown with standard deviations.

### Cell cycle analysis

Cells were treated with DMSO or dTAG-47 for 3, 6, and 9 days. Before staining, cells in a 10-mm-dish at 80% confluence were pulsed with 20μM 5-bromo-2’-deoxyuridine (BrdU) for 2 hours and fixed overnight with 70% ethanol at 4°C. Cells were stained with fluorescein isothiocyanate (FITC)-conjugated anti-BrdU and counterstained with propidium iodide (PI) before analysis by flow cytometry. All flow cytometry figures were generated using FlowJo software.

### Soft Agar colony formation assay

Cells were treated with DMSO or dTAG-47 for 3, 6, and 9 days, and then were plated in 0.3% agarose medium. Cells were fed with culture media with DMSO/dTAG-47 once per week. After 4 weeks, plates were stained with 0.005% Crystal Violet (Sigma), and colonies were counted using a dissecting microscope.

### Immunofluorescence

Cells were seeded on coverslips and treated with DMSO or dTAG-47 for 3, 6, and 9 days. Cell were then fixed with 3.7% paraformaldehyde in PBS at room temperature for 15 minutes. The coverslips were then washed three times with phosphate buffered saline (PBS) and the cells were permeabilized using 0.3% Triton X-100 in PBS at room temperature for 30 minutes. In a humidified chamber, cells were blocked by adding 1% serum in PBS for 30 minutes, and then incubated with primary antibodies (Myogenin, Abcam, ab1835; Myosin Heavy Chain, R&D, MAB4470; HA, Cell Signaling, (C29F4) #3724) with appropriate dilution in 0.5% NP-40 and 1% serum in PBS at 37 °C for 1 hour. After washing 3 times with PBS, cells were incubated in Alexa Fluor secondary antibody (Abcam ab150117, Invitrogen A-11034) with Phalloidin (Thermo Fisher Scientific, A12380) and DAPI diluted in 0.5% NP-40, and 1% serum in PBS at 37 °C for 45 minutes. Coverslips were mounted on slides with Prolong Gold Antifade reagent (Thermo Fisher Scientific, P36930) and dried overnight in the dark. Images were collected using a Nikon fluorescent microscope.

### Nuclei isolation

30 million Rh30 cells were treated with DMSO or dTAG-47 and collected at indicated time points. Cells were washed with ice cold PBS and lysed with cell lysis buffer (10mM Tris-Cl pH7.4, 300mM sucrose, 3mM CaCl_2_, 2mM MgCl_2_, 0.5% NP-40, 5mM DTT, 1mM PMSF, EDTA free protease cocktail inhibitor tablet) using dounce homogenization, nuclei were pelleted by centrifugation and washed with nuclei storage buffer (50mM Tris-Cl pH8.3, 40% glycerol, 5mM MgCl2, 5mM DTT, 0.1 mM EDTA, 1mM PMSF, EDTA free protease cocktail inhibitor tablet). After counting, pelleted nuclei were resuspended in storage buffer, and stored at -80 °C.

### Precision nuclear run-on and sequencing (PRO-seq)

PRO-seq was performed in biological replicates as previously described using approximately 20 million nuclei per run on with GTP, ATP, UTP, and biotin-11-CTP (PerkinElmer) using 0.5% Sarkosyl (Fisher Scientific) to prevent transcription initiation^18,42^. RNA was reversed transcribed and amplified to make the cDNA library for sequencing by the Vanderbilt University Medical Center (VUMC) VANTAGE Genome Sciences Shared Resource on an Illumina Nextseq 500 (SR-75, 50 million reads). The sequences were aligned using bowtie2 (v2.2.2) before using the Nascent RNA Sequencing Analysis (NRSA) pipeline^19^ to determine the gene body and eRNA changes.

### RNA sequencing and data processing

All RNA-seq experiments were performed in biological duplicate. Total RNA was extracted using TRIzol. RNA was submitted to the VUMC VANTAGE core for library preparation and sequencing (Illumima NovaSeq, PE-150, 30 million reads). Adaptors were trimmed using Trimmomatic-0.32 and aligned to the human genome (hg19) using TopHat (v. 2.0.11)^43^. Differential analysis was performed using CuffDiff (v. 2.1.1)^44^.

### Cleavage under targets and release using nuclease (CUT&RUN)

CUT&RUN experiments were performed in biological duplicate as described^16^. Briefly, cells were treated with DMSO or dTAG-47 for the indicated times and 250,000 cells were collected and incubated with Concanavalin A-coated beads (Bangs Laboratories, BP531) for 10 minutes at room temperature, and i with 0.01% freshly dissolved digitonin. Anti-HA (Cell Signaling, (C29F4) #3724), anti-BRD4 (BETHYL, #A301-985A50), anti-H3K4me3 (Abcam, ab12209), anti-MYOD (Santa Cruz, sc-377460), anti-RUNX1 (Santa Cruz, sc-365644), anti-HEB (Santa Cruz, sc-357), anti-ARID1A (Cell Signaling, (D2A8U) #12354), anti-SPT16 (Cell Signaling, D7I2K #12191), anti-CDK8 (Santa Cruz, sc-13155), and anti-PAX3 (Abcam, ab69856) primary antibodies were added and incubated at 4 °C overnight, before washing and binding of secondary antibody (anti-Rabbit, #ab31238, anti-Mouse, #ab46540) for 1hr. After washing, CUTANA pAG-MNase (EpiCypher, #15-1116) fusion protein was added and incubated at 0 °C for 60-90 mins to digest targeted regions of the genome. DNA was then extracted using phenol-chloroform^16^ and sequencing libraries were generated using the NEBNext Ultra II DNA Library Prep Kit for Illumina (NEB #E7645S/L). Sequencing was performed by the VUMC VANTAGE core Illumina NovaSeq (PE-150, 10 million reads).

### Chromatin Immunoprecipitation Sequencing (ChIP-seq)

Cells were treated with dTAG-47 for 0, 2, 4, and 24 hours. Five million cells were used to perform anti-H3K27ac ChIP-seq with *Drosophila* S2 cell spike-in. Cells were cross-linked with 1% formaldehyde for 8 minutes and quenched with 125mM Glycine. Following nuclei isolation, chromatin fragments within 300∼600 bp range were generated by sonication for 25 cycles (30s-on, 30s-off) for 25 cycles with a Biorupter (Diagenode). Chromatin fragments were immunoprecipitated with anti-H3K27ac (Abcam, #ab4729) plus Protein A beads. NEBNext Ultra II DNA Library Prep Kit for Illumina was used to make the DNA libraries (NEB, #E7645S/L), which were sequenced on the Illumina NovaSeq (PE-150, 30 million reads) at the VUMC VANTAGE Shared Resource.

### CUT&RUN and ChIP-seq data analysis

Raw FASTQ data were trimmed using Trimmomatic (v0.32) and paired end reads were aligned to a concatenated human and *E*.*coli* genome (hg19 and ecK12MG1655) for CUT&RUN, or *Drosophila* genome (hg19 and dm3) genome for ChIP-seq using Bowtie2 in very sensitive local mode (--local --very-sensitive-local --no-unal --no-mixed -- no-discordant --phred33 -I 10 -X 700)^45^. Peaks were called using MACS2 with a threshold of q<0.01. Peaks were annotated to the nearest TSS using HOMER. Differential analysis was performed using DiffBind and DESeq2. Significantly changed peaks were defined by a 1.5-fold change threshold and FDR<0.05. BigWig files were generated and normalized with the DESeq2 size factors using Deeptools. Heatmaps were created by Deeptools using the DESeq2 size factor normalized bigwig files.

### Assay for Transposase Accessible Chromatin using sequencing (ATAC-seq)^33^

ATAC-seq was performed in biological duplicate using the ATAC-Seq Kit (Active Motif, catalog No.53150). Briefly, 100,000 Rh30 cells and 2,000 *Drosophila* S2 cells were used to isolate nuclei. Nuclei were incubated with tagmentation Master Mix at 37 °C for 30 minutes after lysing the cells in ice-cold ATAC Lysis Buffer and the DNA was purified with the DNA purification column. PCR amplification of tagmented DNA was performed to make libraries with the appropriate indexed primers. After SPRI bead clean-up, the DNA libraries were sequenced on the Illumina NovaSeq (PE-150, 50 million reads) at the VUMC VANTAGE core.

### ATAC-seq data analysis

Raw FASTQ data were trimmed by Trimmomatic (v0.32), and paired end reads were aligned to a concatenated human and *Drosophila* genome (hg19 and dm3) using Bowtie2 (-X 2000 -q --no-mixed --no-discordant). Peaks were called using Genrich (v. 0.6.1) (https://github.com/jsh58/Genrich) with the following options -j -y -r -e chrM -q 0.01 -a 20. Peaks were annotated to the nearest TSS using HOMER. Differential analysis was performed using DiffBind and DESeq2. Significantly changed peaks were defined by a 1.5-fold change threshold and FDR<0.05. BigWig files were generated and normalized using the DESeq2 size factors using Deeptools. Heatmaps were created by Deeptools using the DESeq2 size factor normalized bigwig files.

### APEX2 in-cell biotinylation

APEX2 in-cell biotinylation was performed as described^46^. Briefly, 100 million cells were incubated with 500 μm biotin phenol (Iris, LS-3500, dissolved in DMSO) at 37 °C for 1 hour. Hydrogen peroxide (H_2_O_2_) was added for 1 minute. After quenching, nuclei were isolated, lysed, and nuclear protein was harvested in RIPA buffer. Protein concentration was determined using DC Protein Assay (BioRad). Biotinylated proteins were purified with streptavidin beads (Thermo Fisher Scientific, #88817) and eluted by boiling in Laemmli sample buffer. Parental Rh30 cells were used as negative control to determine proteins specifically identified in Rh30-PAX3-FOXO1-APEX2 samples. Proteins from parental Rh30 cells and Rh30-PAX3-FOXO1-APEX2 cells were analyzed as biological triplicates that were processed independently.

### FLAG affinity purification

Nuclei were isolated from 100 million cells per sample. The nuclei were then “extracted” by incubating with Pierce Universal Nuclease (Thermo Fisher Scientific, #88701) in co-IP buffer (20 mM Tris pH 8, 150 mM NaCl, 2 mM MgCl_2_, 0.1% NP-40, with protease inhibitors) on ice for 1 hour. Cleared lysates were passed over a 0.45 um cellulose acetate column (Corning) to remove any remaining particulates. Samples were incubated with 60 μl of equilibrated EZ Red Flag M2 bead slurry (EZview by Millipore Sigma, F2426) at 4°C for 2hr in co-IP buffer, After washing, purified proteins were eluted twice with 50 ul of 1 mg/ml 3X flag peptide (Sigma) in co-IP buffer on the nutator for 10 minutes at room temperature.

### Mass Spectrometry

Eluents from Apex or FLAG purifications were prepared for analysis via S-trap trypsin digests using manufacturer’s protocol (S-Trap™ – ProtiFi). The peptides were separated on a self-packed 100 μm × 20 cm reversed phase (Phenomonex - Jupiter 3 micron, 300A) column from which peptides were ionized directly via nano-electrospray into an Exploris 480 (Thermo-Fisher) mass spectrometer. Both full-scan and peptide fragmentation (MS/MS) were collected over the course of a 70-minute aqueous to organic gradient elution in a data-dependent manner using dynamic exclusion to reduce redundancy of peptide acquisition. Resulting MS/MS spectra were searched using SEQUEST^47^ against a human database containing common contaminants and reversed copies of each entry. Resulting identifications were filtered to a 5% false-discovery threshold, collated back to the protein level, and compared across samples using Scaffold (Proteome Software). Filtered total spectral count values were used for fold-change comparisons and p-value estimations.

## Figure Legends

**Extended Figure 1. Analysis of PAX3-FOXO1-FKBP single cell clones. a**, Model of the plasmid DNA template used to insert FKBP12^F36V^-2XHA-P2A-mCherry into the endogenous *PAX3-FOXO1* allele (upper panel). Lower panel shows the PROTAC, dTAG-47, binding to the FKBP12^F36V^ module to link PAX3-FOXO1-FKBP12^F36V^-2xHA to the CRBN E3 ubiquitin ligase to cause rapid degradation of PAX3-FOXO1-FKBP. **b**, Western blot analysis of four Rh30_PAX3-FOXO1-FKBP clones. **c**, Cluster heatmap of Pearson correlations from RNA-seq of four different PAX3-FOXO-FKBP cell lines expanded from single cell clones and parental Rh30 cells. Each cell line has two biological replicates and data was normalized to total counts. **d**, Heatmap of log2 normalized counts from RNA-seq across the four different PAX3-FOXO-FKBP clonal cell lines and parental Rh30 cell line using all the expressed genes; n=2. **e**, Four Rh30_PAX3-FOXO1-FKBP clones were treated with 500 nM dTAG-47, and cell counts were determined using Trypan Blue dye exclusion. Data are mean ± STD (n=3).

**Extended Figure 2. Degradation of PAX3-FOXO1-FKBP triggers cell death and differentiation. a**, Cell cycle analysis of Rh30_PAX3-FOXO1-FKBP cells. The cells were treated with 500 nM dTAG-47 for the indicated times before flow cytometry analysis for BrdU incorporation. Plots of BrdU versus propidium iodide (PI) shows fewer cells in S-phase and accumulation of cells in G1-phase after dTAG-47 treatment. **b**, Bar graph showing statistical analysis of biological replicates of cells in S phase from panel **a**. Data are presented as mean ± STD (n=3); (*p*, independent T test; *: *p* <= 5.0e-02, **: *p* <= 1.0e-02, ***: *p* <= 1.0e-03, ****: *p* <= 1.0e-04). **c**, Growth in soft agar. Rh30_PAX3-FOXO1-FKBP cells were pre-treated with 500 nM dTAG-47 for 6 days before plating in soft agar. 4 weeks later the number of colonies were counted using microscopy. Representative images show colonies using an inverted microscope (10X). The bar graph displays colony counts with the bar the mean ± STD (n=9). (*p*, independent T test. *: *p* <= 5.0e-02, **: *p* <= 1.e-02, ***: *p* <= 1.0e-03, ****: *p* <= 1.0e-04). **d**, Immunofluorescence staining of Myosin Heavy Chain (upper panel) and Myogenin (lower panel). Rh30_PAX3-FOXO1-FKBP cells were treated with 500 nM dTAG-47 for 6 days. DAPI was used to label nuclei (blue). Alexa 568-labeled Phalloidin was used to mark actin filaments (red). Alexa 488 secondary antibody was used to visualize the primary antibody against the skeletal muscle differentiation markers Myosin Heavy Chain and Myogenin (green; 20X).

**Extended Figure 3. Analysis of FOXO1-FKBP12 cells. a**, Western blot analysis of FOXO1-FKBP12 before and after degrading endogenous FOXO1-FKBP12 (upper panel). Lamin B was used as a loading control (bottom panel). **b**, Cell proliferation of Rh30_FOXO1-FKBP12 cells was performed by treating with 500 nM dTAG-47, and cell counts were determined using Trypan Blue dye exclusion. Data are graphed as mean ± STD (n=3). **c**, Flow cytometry analysis of incorporated BrdU versus PI in Rh30-FOXO1-FKBP12 cells after treatment of 500 nM dTAG-47 at Day-3, Day-6 and Day-9. **d**, Bar graph showing the results of biological triplicate flow cytometry analysis showing no change in the percentage of cells in S-phase. Data are presented as mean ± STD (n=3; *p* derived from an independent T test. *: *p* <= 5.0e-02, **: *p* <= 1.0e-02, ***: *p* <= 1.0e-03, ****: *p* <= 1.0e-04). **e**, Immunofluorescence staining of FOXO1 using anti-HA in Rh30_FOXO1-FKBP12 cells. DAPI was used to label nuclei (blue). Alexa 568-labeled Phalloidin was used to mark actin filaments (red). Alexa 488 secondary antibody was used to visualize the primary antibody against HA (green; magnification 100X). **f**, Bar graph showing the number of genes changed detected by RNA-seq after degrading FOXO1 (left panel) or PAX3-FOXO1 (right panel) at indicated time points.

**Extended Figure 4. Gene expression and genomic analysis of PAX3-FOXO1-FKBP cells. a**, Histograms of all H3K4me3 CUT&RUN peaks (upper panel) and the H3K4me3 peaks annotated to genes showing transcriptional changes (lower panel) ±3Kb around the transcription start site (TSS). **b**, Heatmap of RNA-seq plotted using the 717 genes changed after 6hr of dTAG-47 treatment, and plotted using the indicated time points. **c**, Heatmaps plotted using log_2_ transformed fold change (log_2_FC) values of read counts in 200 bp bins ± 5 Kb around the TSSs of the genes with gene body change in at least one time point after degradation of endogenous PAX3-FOXO1. **d**, Heatmap of Log_2_ transformed fold change (log_2_FC) values of pausing indices of all the genes with changes in transcription in at least one time point. **e**, Box plots displays the Log_2_ transformed fold change (log_2_FC) values of PRO-seq at indicated time points (Mann-Whitney U test, *: *p* <= 5.0e-02, **: *p* <= 1.0e-02, ***: *p* <= 1.0e-03, ****: *p* <= 1.00e-04). **f**, Pie chart showing annotation of PAX3-FOXO1 peaks with genomic features using HOMER annotatePeak.pl indicated that most PAX3-FOXO1 binding sites are intergenic or intronic regulatory elements. **g, h**, MA-plots of H3K27ac (**g**) and BRD4 (**h**) peak changes from 2hr to 24hr after degrading PAX3-FOXO1.

**Extended Figure 5. Examples of the super enhancer analysis of PAX3-FOXO1-FKBP cells**. IGV gene tracks showing the PAX3-FOXO1 CUT&RUN, PRO-seq, ATAC-seq, H3K27ac ChIP-seq, and BRD4 CUT&RUN at the super-enhancers associated with the *MYOG* (**a**), *MYCN* (**b**), *MYOD1* (**c**), and *FGGY* loci (**d**).

**Extended Figure 6. Proteomic analysis of PAX3-FOXO1-3xFLAG cells. a**, Motif analysis (de novo) of transcription factors predicted to reside under PAX3-FOXO1 genomic peaks. **b**, Volcano plot generated from the mass spectrometry results of PAX3-FOXO1-3xFLAG tag specific co-IP mass spectrometry. Proteins that purified with FLAG-M2 beads were plotted as log2 fold change (PAX3-FOXO1-3xFLAG/Parental) vs. -log10 of the p-value. The absolute 1.5-fold change and p-value 0.05 was used as the threshold (n = 3 biological replicates, *p* calculated using one-tail unpaired t-test). Significant hits are depicted in blue and red to reflect proteins that are enriched in parental samples and PAX3-FOXO1-APEX2 samples, respectively. **c**, Heatmaps of selected PAX3-FOXO1-associated proteins from the 3xFLAG analysis. Spectral counts are shown within each box. **d**, Venn diagrams displays the overlap of MACS2 identified peaks from CUT&RUN analysis between PAX3-FOXO1 and the indicated factor. **e**, Heatmaps of CUT&RUN signal of the indicated factors around all PAX3-FOXO1 peaks ±3Kb.

**Extended Figure 7. Intersection of CUT&RUN, PRO-seq and ATAC-seq analysis. a**, Venn diagrams showing the overlap between PAX3-FOXO1 peaks and the ATAC-seq peaks. **b**, Heatmaps of ATAC-seq signal over all PAX3-FOXO1 peaks in 10 bp bins ± 3 Kb from the peak center over the 24hr time course after degrading PAX3-FOXO1. **c**, Heatmap of the ATAC-seq signal at the 810 down-regulated sites over the 24hr time course after degrading PAX3-FOXO1. **d**, Motif analysis (de novo) of transcription factors predicted to reside under the ATAC-seq peaks that were changed upon PAX3-FOXO1 degradation. **e**, Pie chart of annotation of the significantly down-regulated ATAC-seq peaks at 2hr after PAX3-FOXO1 degradation to genomic features using HOMER annotatePeak.pl. **f**, Venn diagrams showing the overlap between down-regulated eRNA peaks and the down-regulated ATAC-seq peaks. **g**, Average ATAC-seq signal after degrading PAX3-FOXO1 relative to the center of the significantly down-regulated eRNA peaks. **h**, Heatmaps of CUT&RUN signal of the indicated factors around the ATAC-seq down-regulated peaks (2hr). **i**, Venn diagrams showing the overlap of PAX3-FOXO1 peaks, PAX3 peaks, and down-regulated ATAC-seq peaks. **j**, Heatmap of significant changes in chromatin accessibility at ATAC-seq peaks after a time course of PAX3-FOXO1 degradation. Heatmap is plotted using the peaks significantly changed after a 2hr dTAG-47 treatment. **k**, Box plots displays the Log_2_ transformed fold change (log_2_FC) values of PRO-seq at indicated time points (Mann-Whitney U test, *: *p* <= 5.0e-02, **: *p* <= 1.0e-02, ***: *p* <= 1.0e-03, ****: *p* <= 1.00e-04). **l**, Average ATAC-seq signal after degrading PAX3-FOXO1 relative to the center of the significantly down-regulated eRNA peaks (upper). Histogram of the PRO-seq signal around the significantly down-regulated ATAC-seq peaks (lower). **m**, IGV gene tracks display the change of ATAC-seq and PRO-seq peaks at PAX3-FOXO1-regulated *RUNX2* enhancers over a short time course of degradation of PAX3-FOXO1. Shaded box shows the regulate enhancer element.

**Extended Figure 8. 13 parameter analysis of the KLF4 enhancer cluster after degradation of PAX3-FOXO1-FKBP cells**. IGV gene tracks showing the PAX3-FOXO1-regulated *KLF4* locus and its associated super-enhancer.

**Extended Figure 9. 13 parameter analysis of the PRDM12 enhancer cluster after degradation of PAX3-FOXO1-FKBP cells**. IGV gene tracks showing the PAX3-FOXO1-regulated *PRDM12* demonstrating how some PAX3-FOXO1 bound factors are lost from a specific enhancer.

