## Supplemental figures for "PAX3-FOXO1 coordinates enhancer architecture, eRNA transcription, and RNA polymerase pause release at select gene targets"

Extended data Fig. 1

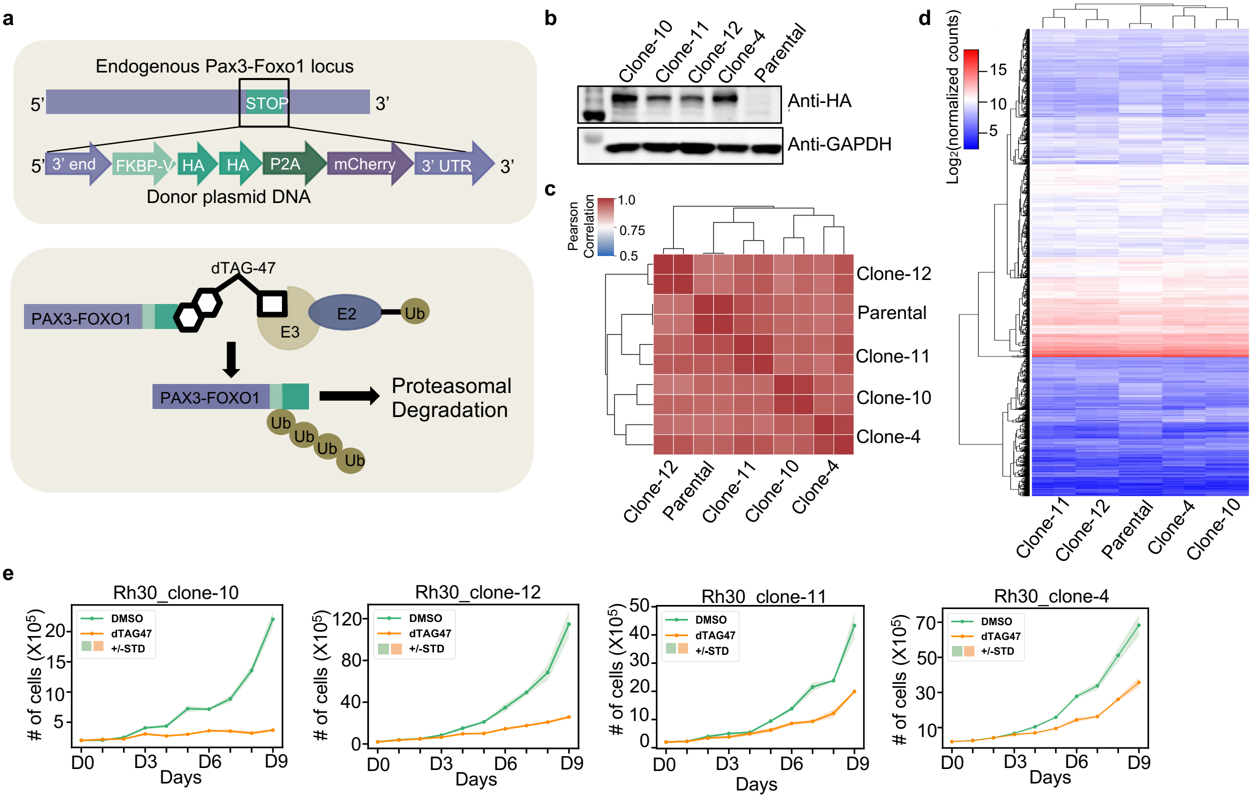


**Extended Figure 1. Analysis of PAX3-FOXO1-FKBP single cell clones. a,** Model of the plasmid DNA template used to insert FKBP12$^{F36V}$-2XHA-P2A-mCherry into the endogenous *PAX3-FOXO1* allele (upper panel). Model showing the PROTAC, dTAG-47, binding to the FKBP12$^{F36V}$ module to link PAX3-FOXO1-FKBP12$^{F36V}$-2xHA to the CRBN E3 ubiquitin ligase to cause rapid degradation of PAX3-FOXO1-FKBP. **b,** Western blot analysis of four Rh30_PAX3-FOXO1-FKBP clones. **c,** Cluster heatmap of Pearson correlations from RNA-seq of four different PAX3-FOXO-FKBP cell lines expanded from single cell clones and parental Rh30 cells. Each cell line has two biological replicates and data was normalized to total counts. **d,** Heatmap of $\log_2$ normalized counts from RNA-seq across the four different PAX3-FOXO-FKBP clonal cell lines and parental Rh30 cell line using all the expressed genes; n=2. **e,** Four Rh30_PAX3-FOXO1-FKBP clones were treated with 500 nM dTAG-47, and cell counts were determined using Trypan Blue dye exclusion. Data are mean ± STD (n=3).

Extended data Fig. 2

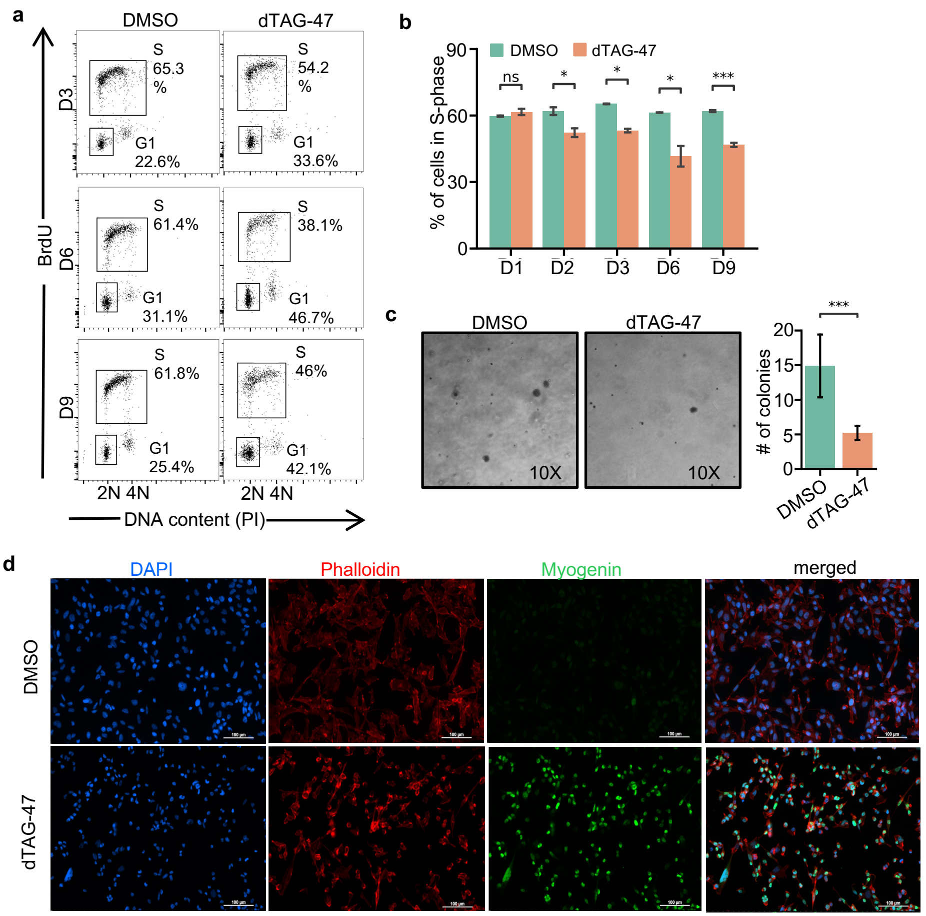


**Extended Figure 2. Degradation of PAX3-FOXO1-FKBP triggers cell death and differentiation. a,** Cell cycle analysis of Rh30_PAX3-FOXO1-FKBP cells. The cells were treated with 500 nM dTAG-47 for the indicated times before flow cytometry analysis for BrdU incorporation. Plots of BrdU versus propidium iodide (PI) shows fewer cells in S-phase and accumulation of cells in $G_1$-phase after dTAG-47 treatment. **b,** Bar graph showing statistical analysis of biological replicates of cells in S phase from panel a. Data are presented as mean ± STD (n=3); (*p*, independent T test; *: *p* <= 5.0e-02, **: *p* <= 1.0e-02, ***: *p* <= 1.0e-03, ****: *p* <= 1.0e-04). **c,** Growth in soft agar. Rh30_PAX3-FOXO1-FKBP cells were pre-treated with 500 nM dTAG-47 for 6 days before plated in soft agar. 4 weeks later the number of colonies were counted using microscopy. Representative images show colonies using an inverted microscope (10X). The bar graph displays colony counts with the bar the mean ± STD (n=9). (*p*, independent T test. *: *p* <= 5.0e-02, **: *p* <= 1.e-02, ***: *p* <= 1.0e-03, ****: p <= 1.0e-04). **d,** Immunofluorescence analysis of Myogenin expression. Rh30_PAX3-FOXO1-FKBP cells were treated with 500 nM dTAG-47 for 6 days. DAPI was used to label nuclei (blue). Alexa 568-labeled Phalloidin was used to mark actin filaments (red). Alexa 488 secondary antibody was used to visualize the primary antibody against skeletal muscle differentiation markers Myosin Heavy Chain and Myogenin (green; 20X).

Extended data Fig. 3

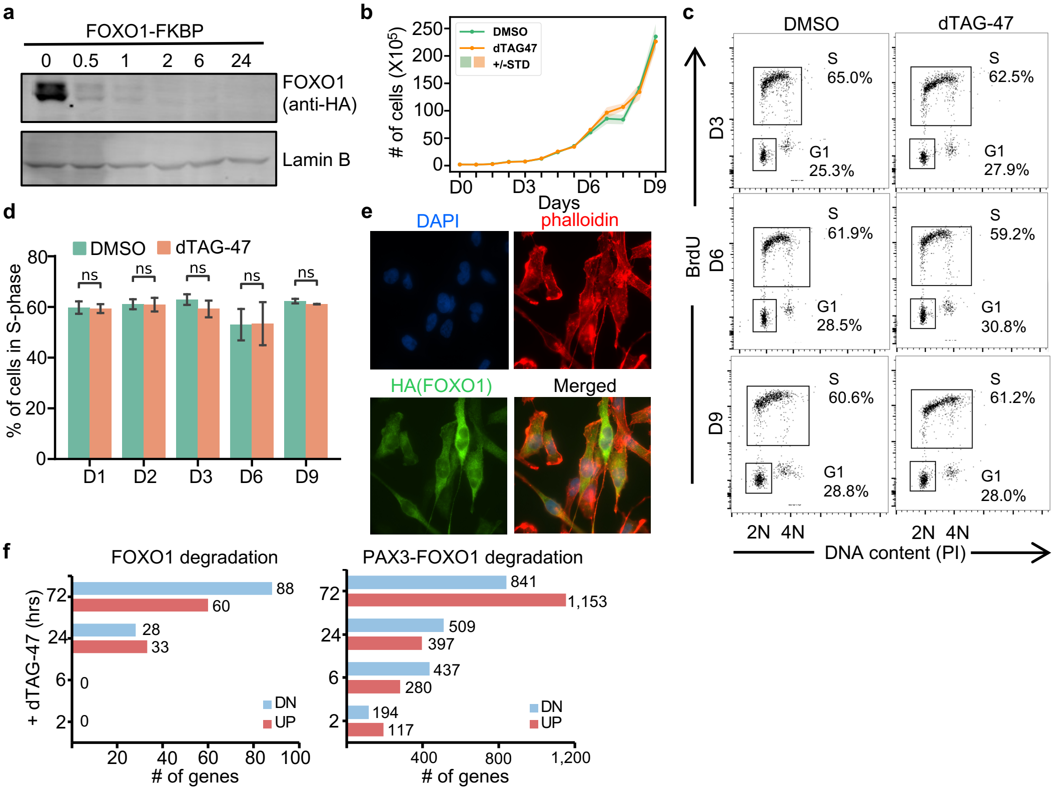


**Extended Figure 3. Analysis of FOXO1-FKBP12 cells**. **a,** Western blot analysis of FOXO1-FKBP12 before and after degrading endogenous FOXO1-FKBP12 (upper panel). Lamin B was used as a loading control (bottom panel). **b,** Cell proliferation of Rh30_FOXO1-FKBP12 cells was performed by treating with 500 nM dTAG-47, and cell counts were determined using Trypan Blue dye exclusion. Data are graphed as mean ± STD (n=3). **c,** Flow cytometry analysis of incorporated BrdU versus PI in Rh30-FOXO1-FKBP12 cells after treatment of 500 nM dTAG-47 at Day-3, Day-6 and Day-9. **d,** Bar graph showing the results of biological triplicate flow cytometry analysis showing no change in the percentage of cells in S-phase. Data are presented as mean ± STD (n=3; *p* derived from an independent T test. *: *p* <= 5.0e-02, **: *p* <= 1.0e-02, ***: *p* <= 1.0e-03, ****: *p* <= 1.0e-04). **e,** Immunofluorescence staining of FOXO1 using anti-HA in Rh30_FOXO1-FKBP12 cells. DAPI was used to label nuclei (blue). Alexa 568-labeled Phalloidin was used to mark actin filaments (red). Alexa 488 secondary antibody was used to visualize the primary antibody against HA (green; magnification 100X). **f,** Bar graph showing the number of gene changed detected by RNA-seq after degrading FOXO1 (left panel) or PAX3-FOXO1 (right panel) at indicated time points.

**Extended data Fig. 4**

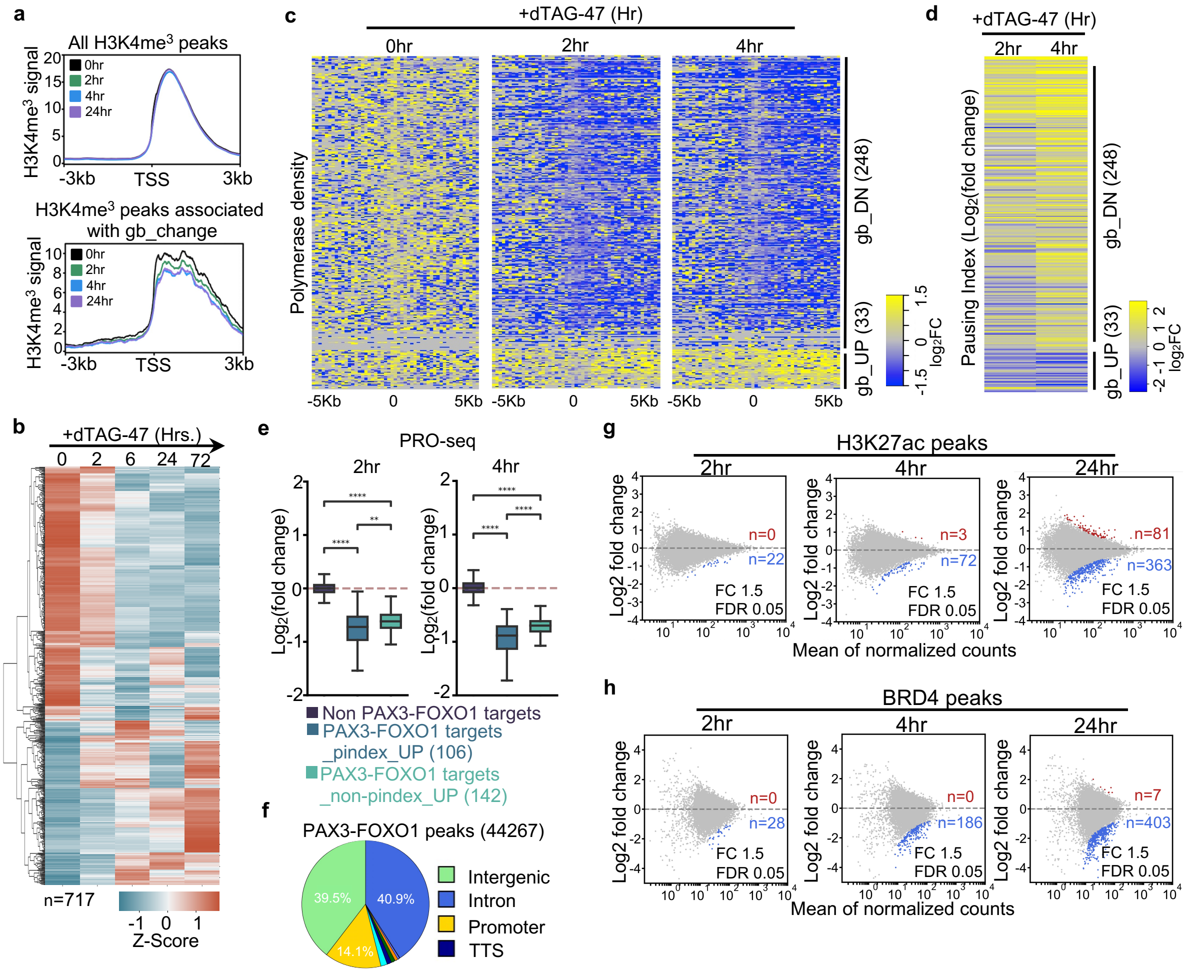


**Extended Figure 4. Gene expression and genomic analysis of PAX3-FOXO1-FKBP cells**. **a,** Histograms of all H3K4me3 CUT&RUN peaks (upper panel) and the H3K4me3 peaks annotated to genes showing transcriptional changes (lower panel) $\pm$3Kb around the transcription start site (TSS). **b,** Heatmap of RNA-seq plotted by the genes changed after 6hr of dTAG-47 treatment at indicated time points. **c,** Heatmaps plotted using $\log_2$ transformed fold change ($\log_2$FC) values of read counts in 200 bp bins $\pm$ 5 Kb around the TSSs of the genes with gene body change in at least one time point after degradation of endogenous PAX3-FOXO1. **d,** Heatmap of $\text{Log}_2$ transformed fold change ($\log_2$FC) values of pausing indices of all the genes with changes in transcription in at least one time point. **e,** Box plots displays the $\text{Log}_2$ transformed fold change ($\log_2$FC) values of PRO-seq at indicated time points (Mann-Whitney U test, *: $p <= 5.0e-02$, **: $p <= 1.0e-02$, ***: $p <= 1.0e-03$, ****: $p <= 1.00e-04$). **f,** Pie chart showing annotation of PAX3-FOXO1 peaks with genomic features using HOMER annotatePeak.pl indicated that most PAX3-FOXO1 binding sites are intergenic or intronic regulatory elements. **g, h,** MA-plots of H3K27ac (**g**) and BRD4 (**h**) peak changes from 2hr to 24hr after degrading PAX3-FOXO1.

**Extended data Fig. 5**

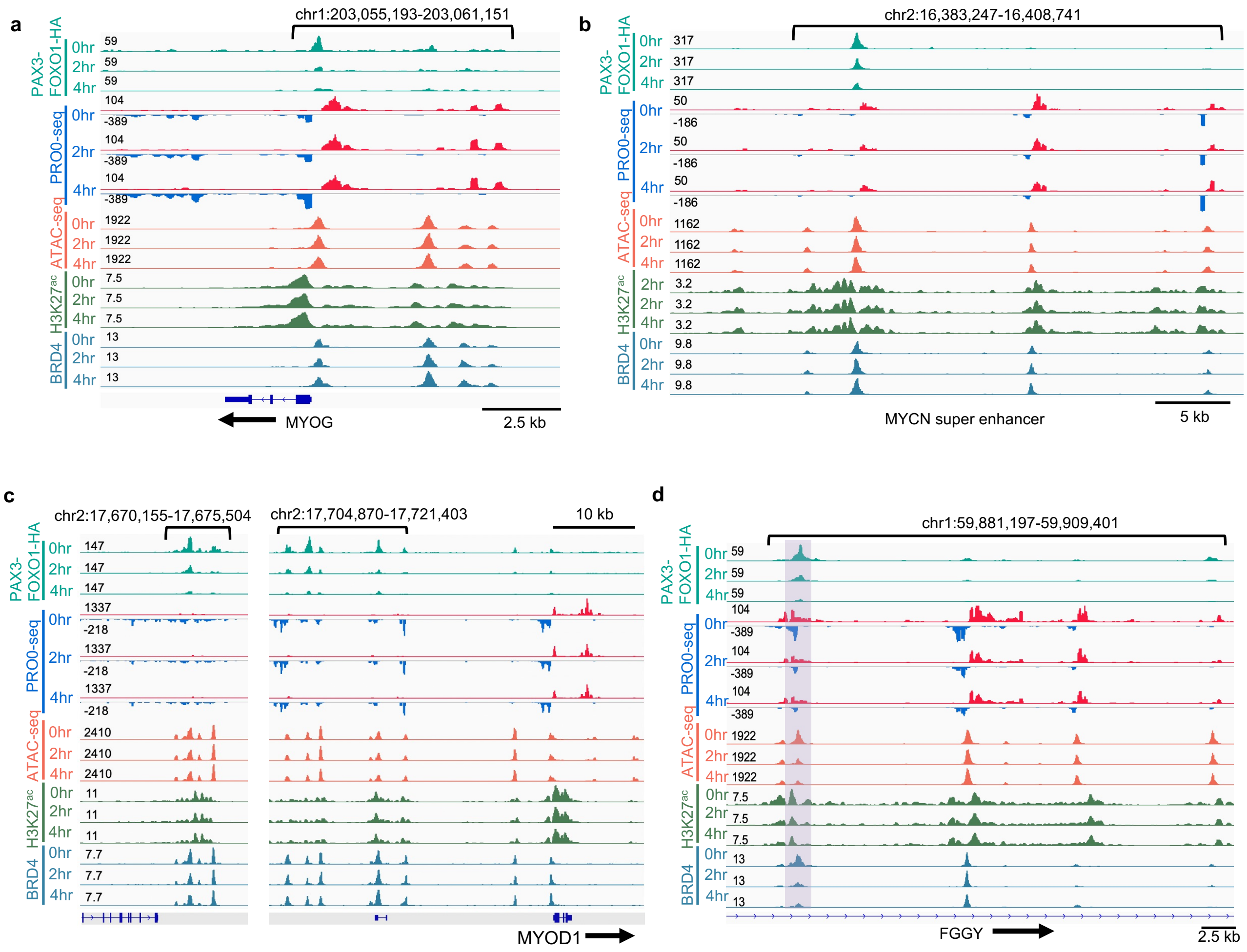


**Extended Figure 5. Examples of the super enhancer analysis of PAX3-FOXO1-FKBP cells.** IGV gene tracks showing the PAX3-FOXO1 CUT&RUN, PRO-seq, ATAC-seq, H3K27ac ChIP-seq, and BRD4 CUT&RUN at the super-enhancers associated with the *MYOG* (**a**), *MYCN* (**b**), *MYOD1* (**c**), and *FGGY* loci (**d**).

**Extended data Fig. 6**

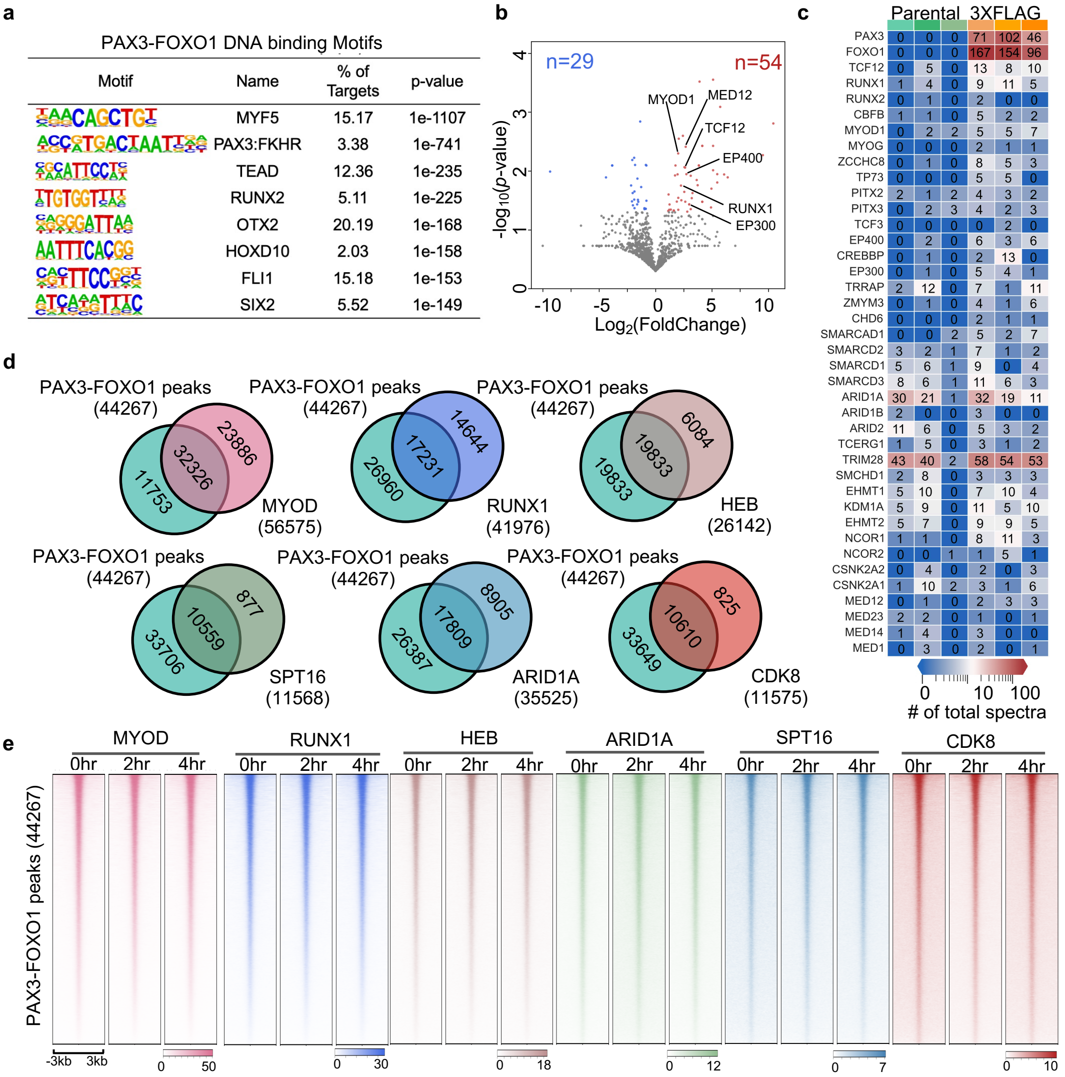


**Extended Figure 6. Proteomic analysis of PAX3-FOXO1-3XFLAG cells. a,** Motif analysis (de novo) of transcription factors predicted to reside under PAX3-FOXO1 genomic peaks. **b,** Volcano plot generated from the mass spectrometry results of PAX3-FOXO1-3XFLAG tag specific co-IP mass spectrometry. Proteins that purified with FLAG-M2 beads were plotted as log2 fold change (PAX3-FOXO1-3xFLAG/Parental) vs. -log10 of the p-value. The absolute 1.5-fold change and p-value 0.05 was used as the threshold (n = 3 biological replicates, *p* calculated using one-tail unpaired T-test). Significant hits are depicted in blue and red to reflect proteins that are enriched in parental samples and PAX3-FOXO1-APEX2 samples, respectively. **c,** Heatmaps of selected PAX3-FOXO1-associated proteins from the 3XFLAG analysis. Spectral counts are shown within each box. **d,** Venn diagrams displays the overlap of MACS2 identified peaks from CUT&RUN analysis between PAX3-FOXO1 and the indicated factor. **e,** Heatmaps of CUT&RUN signal of the indicated factors around all PAX3-FOXO1 peaks ±3Kb.

**Extended data Fig. 7**

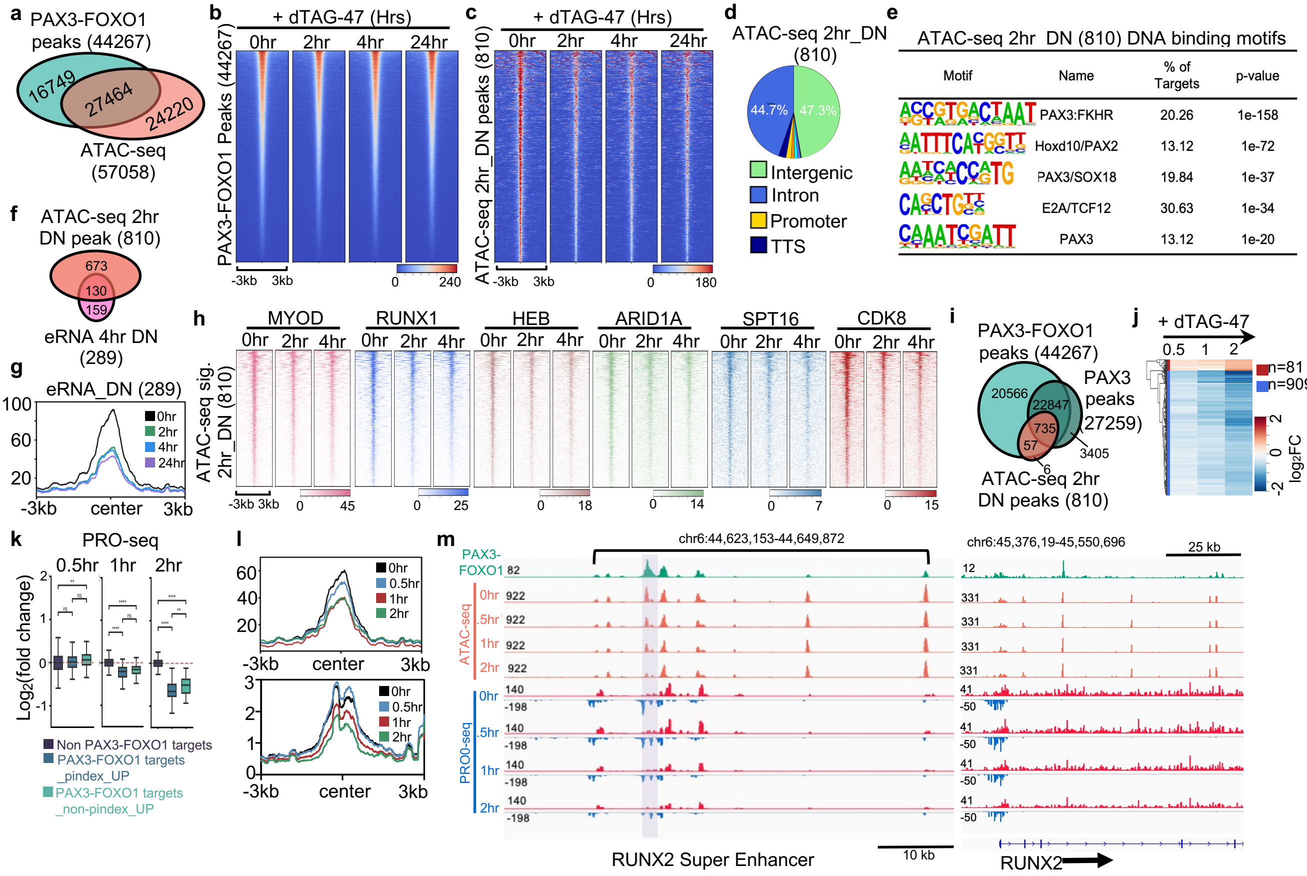


**Extended Figure 7. Intersection of CUT&RUN, PRO-seq and ATAC-seq analysis.** **a,** Venn diagrams showing the overlap between PAX3-FOXO1 peaks and the ATAC-seq peaks. **b,** Heatmaps of ATAC-seq signal over all PAX3-FOXO1 peaks in 10 bp bins ± 3 Kb from the peak center over the 24hr time course after degrading PAX3-FOXO1. **c,** Heatmap of the ATAC-seq signal at the 810 down-regulated sites over the 24hr time course after degrading PAX3-FOXO1. **d,** Pie chart of annotation of the significantly down-regulated ATAC-seq peaks at 2hr after PAX3-FOXO1 degradation to genomic features using HOMER annotatePeak.pl. **e,** Motif analysis (de novo) of transcription factors predicted to reside under the ATAC-seq peaks that were changed upon PAX3-FOXO1 degradation. **f,** Venn diagrams showing the overlap between down-regulated eRNA peaks and the down-regulated ATAC-seq peaks. **g.** Average ATAC-seq signal after degrading PAX3-FOXO1 relative to the center of the significantly down-regulated eRNA peaks. **h,** Heatmaps of CUT&RUN signal of the indicated factors around the ATAC-seq down-regulated peaks (2hr). **i,** Venn diagrams showing the overlap of PAX3-FOXO1 peaks, PAX3 peaks, and down-regulated ATAC-seq peaks. **j,** Heatmap of significant changes in ATAC-seq peaks after a time course of PAX3-FOXO1 degradation. Heatmap is plotted using the peaks significantly changed after a 2hr dTAG-47 treatment. **k,** Box plots display the $Log_2$ transformed fold change ($log_2FC$) values of PRO-seq at indicated time points (Mann-Whitney U test, *: $p <= 5.0e-02$, **: $p <= 1.0e-02$, ***: $p <= 1.0e-03$, ****: $p <= 1.00e-04$). **l,** Average ATAC-seq signal after degrading PAX3-FOXO1 relative to the center of the significantly down-regulated eRNA peaks (upper). Histogram of the PRO-seq signal around the significantly down-regulated ATAC-seq peaks (lower). **m,** IGV gene tracks display the change of ATAC-seq and PRO-seq peaks at PAX3-FOXO1-regulated *RUNX2* enhancers over a short time course of degradation of PAX3-FOXO1. Shaded box shows the regulated enhancer element.

**Extended data Fig. 8**

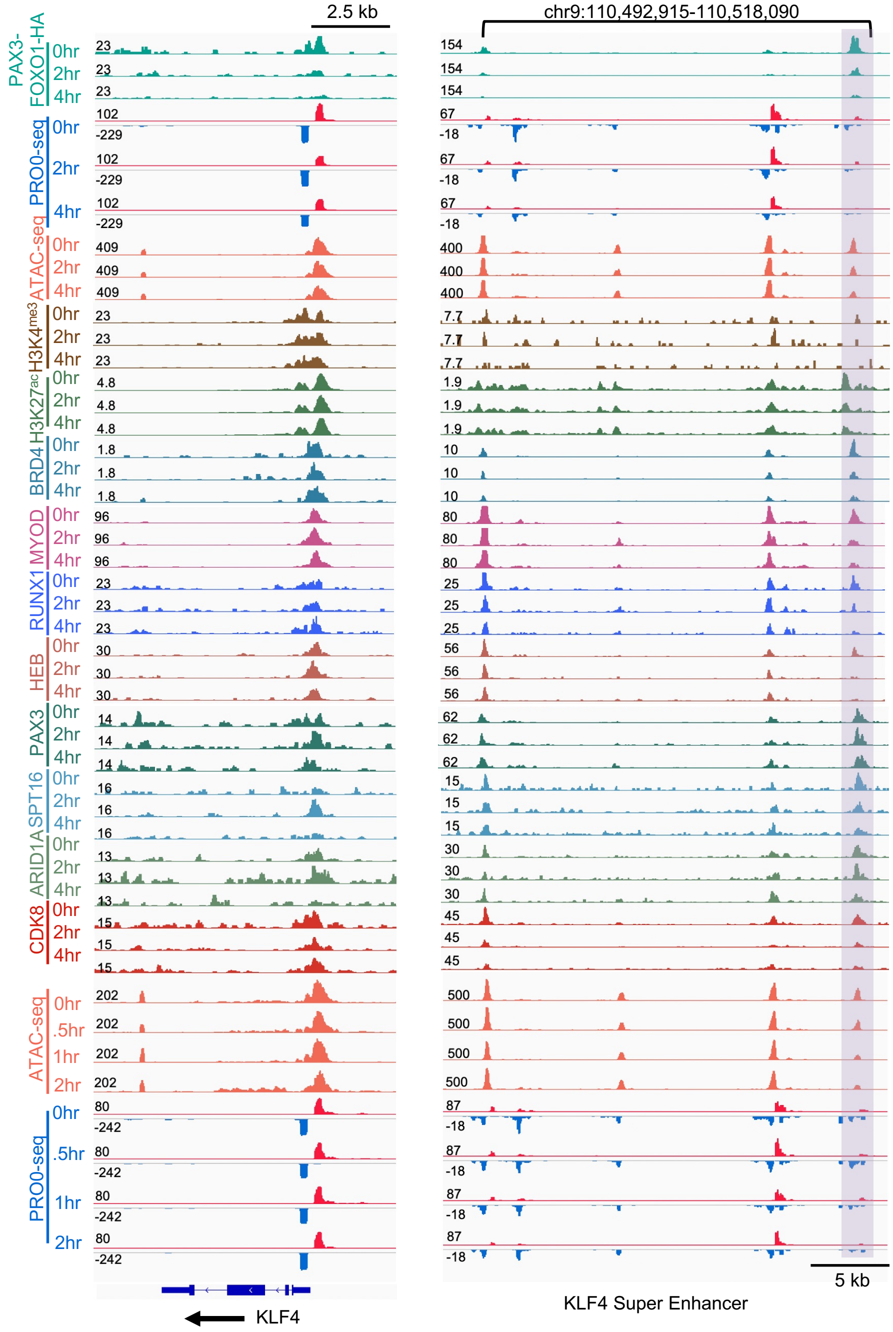


**Extended Figure 8. 13 parameter analysis of the KLF4 enhancer cluster after degradation of PAX3-FOXO1-FKBP cells.** IGV gene tracks showing the PAX3-FOXO1-regulated *KLF4* locus and its associated super-enhancer.

**Extended data Fig. 9**

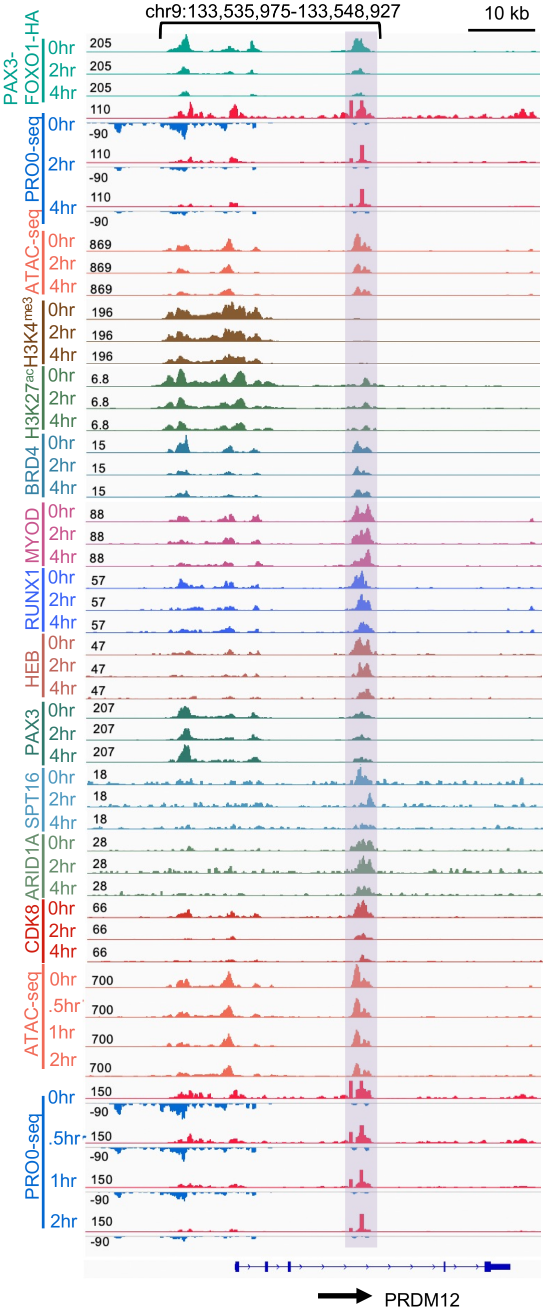


**Extended Figure 9. 13 parameter analysis of the PRDM12 enhancer cluster after degradation of PAX3-FOXO1-FKBP cells.** IGV gene tracks showing the PAX3-FOXO1-regulated *PRDM12* demonstrating how some PAX3-FOXO1 bound factors are lost from a specific enhancer.
